# Kinome-wide CRISPR/Cas9-knockout screening reveals critical protein kinases in vasopressin V2-receptor signaling

**DOI:** 10.64898/2026.07.03.736393

**Authors:** Euijung Park, Lihe Chen, Viswanathan Raghuram, Shaza Khan, Adrian R. Murillo-de-Ozores, Chung-Lin Chou, Chin-Rang Yang, Mark A. Knepper

## Abstract

Identification of signaling networks is an essential goal in systems biology. Here, we use CRISPR/Cas9 knockout screening (employing a whole kinome sgRNA library) to identify functionally critical protein kinases in a well-studied G_α_s-dependent G-protein coupled receptor (GPCR)-signaling model, namely the vasopressin V2 receptor (V2R) pathway. Screening was done using a specially-designed fluorescence-based reporter cell line with green-fluorescent protein (GFP) co-transcribed with *Aqp2*, a gene whose transcription is dependent on vasopressin-mediated activation of protein kinase A (PKA). Positive regulators (n=14) included PKA-catalytic subunit α (Prkaca) and Dyrk1a (*minibrain* homolog). Negative regulators (n=12) included PKA-regulatory subunit type Iα, Stk11 (catalytic subunit of liver kinase B1 [LKB1] complex), and three TGF-β receptor subunits (Tgfbr1, Tgfbr2, Tgfbr3) (see https://esbl.nhlbi.nih.gov/Databases/Kinome-CRISPR-screen/ for full list). Dyrk1a knockout cell lines failed to express AQP2 protein and exhibited a profound decrease in AQP2 mRNA. RNA-sequencing demonstrated widespread increases in cell-cycle transcripts, with a general defect in cell differentiation, accounting for AQP2 loss. TGF-β exposure to un-transformed cells results in a profound decrease in V2R and AQP2 mRNA abundance along with multiple additional transcriptional targets of V2R-PKA signaling, consistent with prior findings in TGF-β-mediated vasopressin ‘escape’. Stk11/LKB1 knockout lines displayed marked increases in AQP2 protein and mRNA, even in the absence of vasopressin. RNA-sequencing showed a marked similarity between the responses to Stk11/LKB1 deletion and vasopressin exposure in untransformed cells. Phospho-proteomic data point to opposing roles of Stk11/LKB1 and PKA in the regulation of cAMP-responsive transcriptional coactivator (CRTC) proteins in the transcriptional response to V2R-PKA signaling.

**Significance Statement:** Cells throughout the body are regulated by extracellular signals like the hormone, vasopressin. Hormonal effects on cellular function are mediated by membrane receptors that trigger biochemical changes, often by inducing chemical modification of the amino acids making up individual proteins, such as addition of function-altering phosphate groups (phosphorylation). Protein phosphorylation is mediated by enzymes known as “protein kinases”. Here, we have screened all known protein kinases using modern CRISPR/Cas9 technology to identify those involved in vasopressin action in the kidney. As expected from prior knowledge, the screen identified protein kinase A and one of its regulatory subunits, but also identified several protein kinases not previously implicated in vasopressin action in the kidney.

## Introduction

Much of intracellular signaling depends on post-translational modifications of proteins, most prominently protein phosphorylation. Consequently, to identify a signaling network, it is necessary to identify the proteins that regulate protein phosphorylation, namely protein kinases and protein phosphatases. The introduction of CRISPR knockout screening approaches (1–4) brings forth a new way to identify functional elements of a given signaling system. Here, we use CRISPR knockout screening, employing a whole-kinome sgRNA library to identify novel critical protein kinases in a previously well-studied G protein coupled receptor (GPCR)-dependent pathway, namely, that of the vasopressin V2 receptor (V2R) (6–7), which plays a key role in the kidney by regulating water excretion (8) largely through control of the aquaporin-2 (AQP2) water channel (9).

The vasopressin V2 receptor signals via G_α_s-mediated activation of the adenylyl cyclase ADCY6, and generation of cyclic AMP (cAMP) (10). It is one of many G_α_s-coupled GPCRs expressed in a variety of tissues (11). Although cAMP can produce cellular responses through several mechanisms, the physiological effects of V2 receptor activation on osmotic water transport appear to be mediated chiefly by protein kinase A (PKA) (12), specifically PKA-catalytic subunit-α (PKA-Cat-α) (13). In the kidney collecting duct, V2 receptor dependent signaling regulates water transport via two chief mechanisms: 1) regulation of membrane trafficking of vesicles containing the aquaporin-2 (AQP2) water channel to and from the plasma membrane (14); and 2) regulation of transcription of the aquaporin-2 gene, *Aqp2* (15–17). Failure of the latter process is the basis of a variety of water-balance disorders in humans (18). Extensive protein mass spectrometry-based quantitative phosphoproteomics studies in kidney collecting duct cells have demonstrated that V2 receptor signaling results in increased phosphorylation at many known PKA target sites in various proteins and has identified many new putative PKA target sites (19–22). However, such studies have also demonstrated many phosphorylation changes at amino acid sites that are not typical of PKA targets, most likely indicating the involvement of additional protein kinases in V2 receptor signaling. Deletion of either PKA-Cat-α alone (13) or together with PKA-Cat-β (12), paints a similar picture with a marked reduction of phosphorylation at many amino acid sites that are typical of PKA signaling, but also with changes in phosphorylation at a large number of sites that are phosphorylated by other yet-to-be-identified protein kinases.

To identify novel protein kinases important to the V2 receptor network and its regulation of *Aqp2* transcription, we have carried out CRISPR/Cas9 screening with a whole-kinome sgRNA library using a specially designed fluorescence-based reporter system with a green-fluorescent protein (GFP) cassette co-transcribed with *Aqp2* in a vasopressin-responsive collecting duct cell line. These cells express the vasopressin V2 receptor endogenously. This screening identified 14 positive regulators and 12 negative regulators of *Aqp2* transcription in the presence of vasopressin. We focused on the strongest positive regulators (Dyrk1a and PKA-Cat-α) and selected protein kinases found to be among the strongest negative regulators (Stk11/LKB1) and all three subunits of the TGF-β receptor-kinase (Tgfbr1, Tgfbr2 and Tgfbr3) for further studies to identify their roles in V2 receptor-mediated regulation of *Aqp2* transcription.

## Results and Discussion

Prior studies have shown that *Aqp2* transcription is almost completely dependent on vasopressin V2 receptor (V2R) signaling through activation of the V2R-G_α_s-ADCY6-cAMP-PKA pathway (16–17). To identify protein kinases involved, we used a CRISPR/Cas9 knockout (KO) screening strategy employing a sgRNA library targeting 714 protein kinases and related proteins (4 sgRNAs per target) (**Figure 1A**) with a custom reporter system in which GFP was co-transcribed with the aquaporin-2 gene (gene symbol: *Aqp2*) in collecting duct mpkCCD cells (**Figure 1B**). These cells are modified from those described in our prior report (23) to constitutively express Cas9. Cells exposed to a V2 receptor selective vasopressin analog (dDAVP) were classified into four groups (‘Neg’, ‘Low’, ‘Mid’, ‘High’ GFP intensity) (**Figure 1C**). With this strategy, a protein kinase “hit” could identify either a kinase directly involved in the V2R signaling pathway, or a kinase involved in more general processes, e.g. regulation of the state of cell differentiation by altering epithelial cell polarity or cell-cycle entry (*vide infra*).

**Figure 1.**
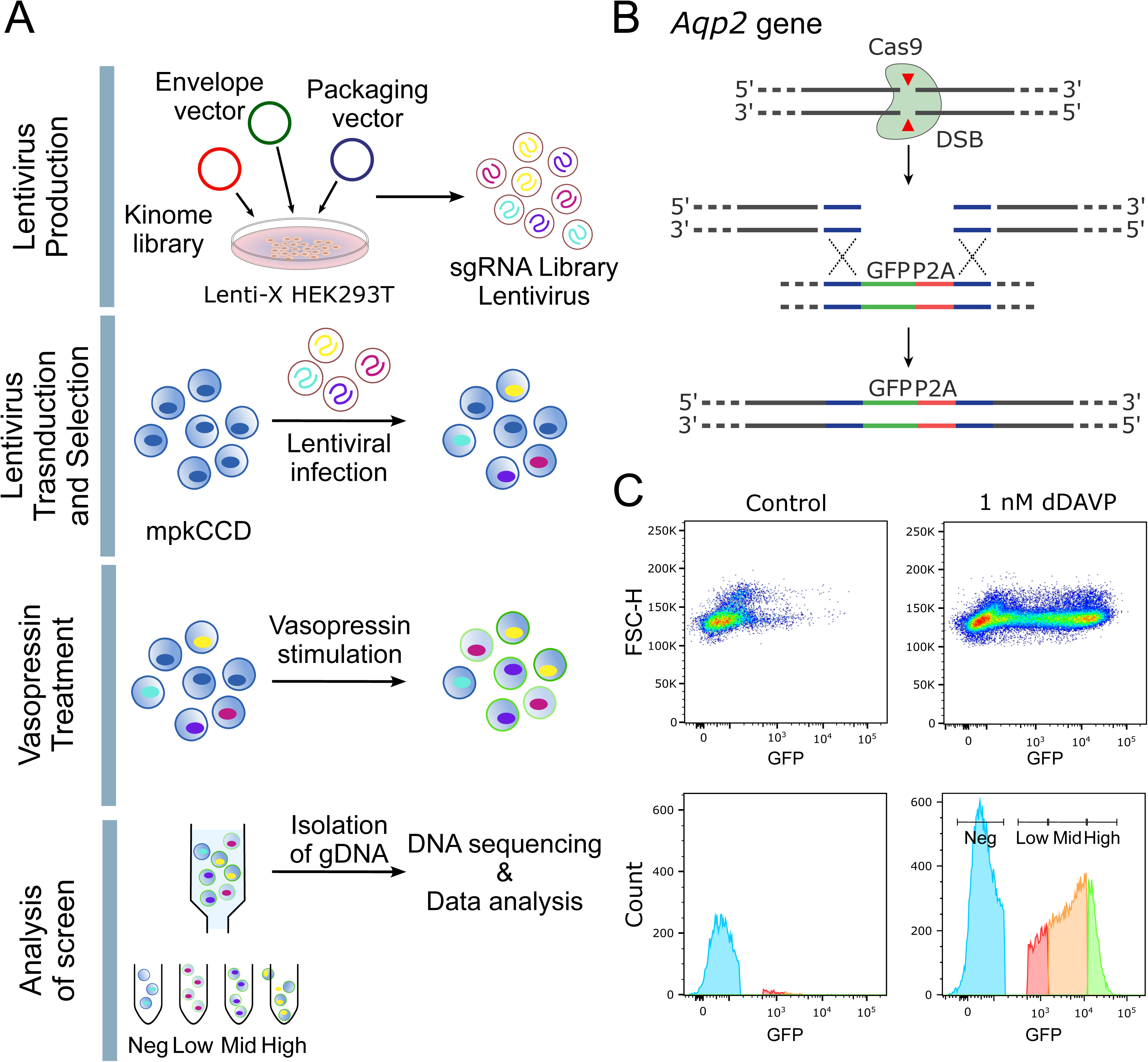
CRISPR kinome screening methodology. **A.** Schematic workflow for CRISPR-mediated kinome screening. **B.** Reporter cell line. GFP cassette was inserted 5’ to *Aqp2* start codon in Cas9-expressing mpkCCD cells, allowing co-transcription of GFP and *Aqp2*. **C.** Cell sorting based on GFP intensity in non-transformed cells shows a marked shift to higher fluorescence levels with dDAVP (1 nM, 48 hours), documenting cellular responsiveness to dDAVP. dDAVP-treated cells were classified into four groups (‘Neg’, ‘Low’, ‘Mid’, ‘High’ GFP intensity).

The kinome CRISPR/Cas9 screen (**Figure 2A**) identified 26 protein kinase proteins that, when deleted, significantly affected GFP fluorescence in the reporter cell line. As expected, Prkaca (PKA-Cat-α) was reported out as a strong positive regulator of *Aqp2* transcription and Prkar1a (PKA regulatory subunit, type Iα) was identified as a strong negative regulator. The latter is an inhibitor of PKA activity that is released from its inhibitory interaction with the catalytic subunit when it binds cAMP. Identification of Prkaca and Prkar1a provides strong confirmation of the validity of the CRISPR screening approach. Beyond this, the screen identified several genes that code for proteins that function together in the same complex (Stk11 and Strada [LKB1 complex]; Tgfbr1, Tgfbr2 and Tgfbr3 [transforming growth factor β receptor complex]) or the same pathway (Map3k1, Map2k4 and Map2k7 [JNK pathway]); Wnk4, Oxsr1 and Ulk1 [Na^+^ transport regulation]. Identification of so many of these protein groups is unlikely as random events and further bolsters confidence in the validity of the overall approach. A full list of the genes identified in the kinome CRISPR KO screen with annotations is provided at https://esbl.nhlbi.nih.gov/Databases/Kinome-CRISPR-screen/ and in abbreviated form in **Table 1**. The CRISPR screen identified several novel putative regulators of *Aqp2* gene transcription, including Dyrk1a (Dual-Specificity Tyrosine Phosphorylation-Regulated Kinase 1a) and Stk11 (Liver Kinase B 1) (**Figure 2B**). **Figure 2C** highlights key CRISPR screening hits that were selected for further study. Among these, Prkaca has already been demonstrated to play an important role in signaling from the vasopressin V2 receptor and other G_α_s-coupled GPCRs. Beyond this, Dyrk1a, Stk11 and Strada can be considered candidates for specialized roles in the V2R-G_α_s-ADCY6-cAMP-PKA pathway. Furthermore, because of the identification of Tgfbr1, Tgfbr2 and Tgfbr3 as putative negative regulators and prior data implicating TGF-β in development of vasopressin-resistance in the syndrome of inappropriate antidiuresis (‘vasopressin escape’) (24), we also investigate actions of TGF-β1 in additional studies.

**Figure 2.**
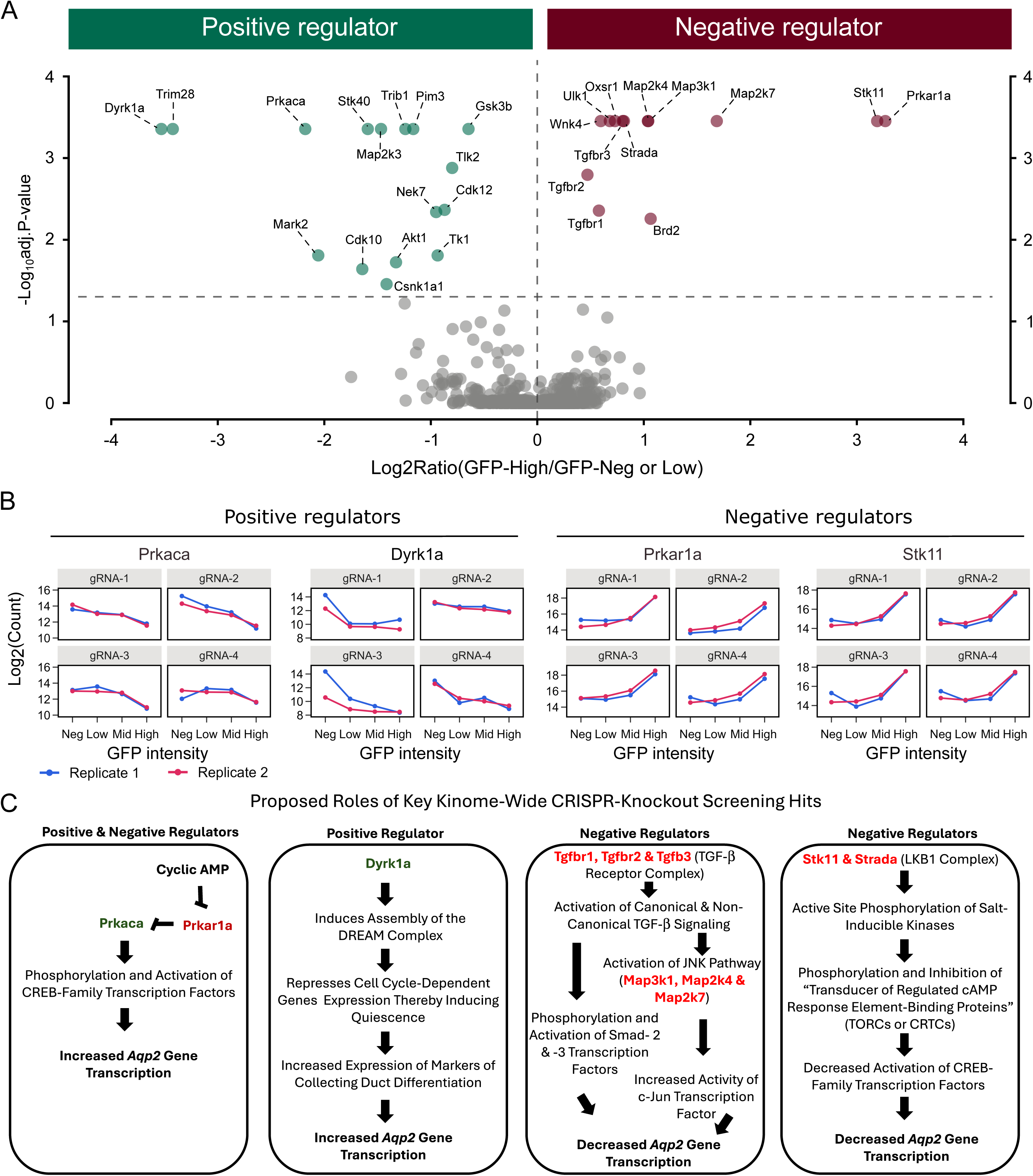
Kinome CRISPR KO screening results. **A.** Volcano plot showing effect of CRISPR KO employing sgRNAs targeting 714 protein kinases and related proteins (n=4 sgRNA per target). Analysis used mean values for two independent replicates for all four sgRNAs per target and is based on normalized sgRNA counts corresponding to “GFP-High” group compared to the combined the “GFP-Neg” and “GFP-Low” groups. **B**. Detailed data for two individual replicates and each of four sgRNAs corresponding to four protein kinase genes: Positive regulators, Prkaca and Dyrk1a; negative regulators, Prkar1a and Stk11. **C.** Proposed roles of key CRISPR KO screening “hits”. Positive regulators are listed in green; negative regulators in red. This panel serves as an hypothesis list for the remainder of this paper.

**Table 1.**
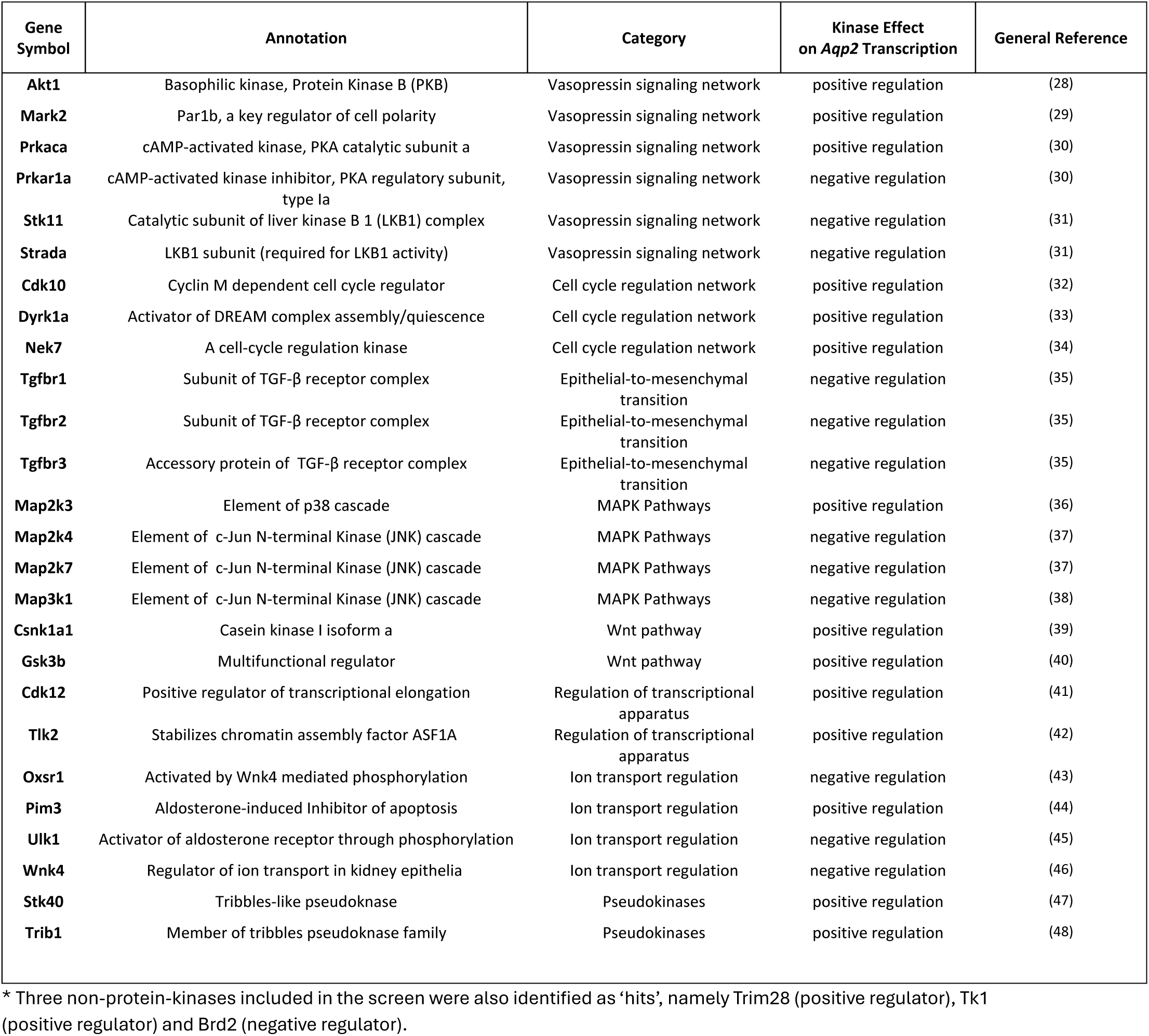
Summary of kinome CRISPR KO screening ‘hits’*.

| Gene Symbol | Annotation | Category | Kinase Effect on *Aqp2* Transcription | General Reference |
| --- | --- | --- | --- | --- |
| **Akt1** | Basophilic kinase, Protein Kinase B (PKB) | Vasopressin signaling network | positive regulation | (28) |
| **Mark2** | Par1b, a key regulator of cell polarity | Vasopressin signaling network | positive regulation | (29) |
| **Prkaca** | cAMP-activated kinase, PKA catalytic subunit a | Vasopressin signaling network | positive regulation | (30) |
| **Prkar1a** | cAMP-activated kinase inhibitor, PKA regulatory subunit, type Ia | Vasopressin signaling network | negative regulation | (30) |
| **Stk11** | Catalytic subunit of liver kinase B 1 (LKB1) complex | Vasopressin signaling network | negative regulation | (31) |
| **Strada** | LKB1 subunit (required for LKB1 activity) | Vasopressin signaling network | negative regulation | (31) |
| **Cdk10** | Cyclin M dependent cell cycle regulator | Cell cycle regulation network | positive regulation | (32) |
| **Dyrk1a** | Activator of DREAM complex assembly/quiescence | Cell cycle regulation network | positive regulation | (33) |
| **Nek7** | A cell-cycle regulation kinase | Cell cycle regulation network | positive regulation | (34) |
| **Tgfb1** | Subunit of TGF-β receptor complex | Epithelial-to-mesenchymal transition | negative regulation | (35) |
| **Tgfb2** | Subunit of TGF-β receptor complex | Epithelial-to-mesenchymal transition | negative regulation | (35) |
| **Tgfb3** | Accessory protein of TGF-β receptor complex | Epithelial-to-mesenchymal transition | negative regulation | (35) |
| **Map2k3** | Element of p38 cascade | MAPK Pathways | positive regulation | (36) |
| **Map2k4** | Element of c-Jun N-terminal Kinase (JNK) cascade | MAPK Pathways | negative regulation | (37) |
| **Map2k7** | Element of c-Jun N-terminal Kinase (JNK) cascade | MAPK Pathways | negative regulation | (37) |
| **Map3k1** | Element of c-Jun N-terminal Kinase (JNK) cascade | MAPK Pathways | negative regulation | (38) |
| **Csnk1a1** | Casein kinase I isoform a | Wnt pathway | positive regulation | (39) |
| **Gsk3b** | Multifunctional regulator | Wnt pathway | positive regulation | (40) |
| **Cdk12** | Positive regulator of transcriptional elongation | Regulation of transcriptional apparatus | positive regulation | (41) |
| **Tlk2** | Stabilizes chromatin assembly factor ASF1A | Regulation of transcriptional apparatus | positive regulation | (42) |
| **Oxsr1** | Activated by Wnk4 mediated phosphorylation | Ion transport regulation | negative regulation | (43) |
| **Pim3** | Aldosterone-induced Inhibitor of apoptosis | Ion transport regulation | positive regulation | (44) |
| **Ulk1** | Activator of aldosterone receptor through phosphorylation | Ion transport regulation | negative regulation | (45) |
| **Wnk4** | Regulator of ion transport in kidney epithelia | Ion transport regulation | negative regulation | (46) |
| **Stk40** | Tribbles-like pseudokinase | Pseudokinases | positive regulation | (47) |
| **Trib1** | Member of tribbles pseudokinase family | Pseudokinases | positive regulation | (48) |

\* Three non-protein-kinases included in the screen were also identified as ‘hits’, namely Trim28 (positive regulator), TK1 (positive regulator) and Brd2 (negative regulator).

### Prkaca and Prkar1a

In the kinome CRISPR screen, PKA catalytic subunit α (PKA-Cat-α) showed up as a positive regulator of *Aqp2* transcription, whereas PKA-Cat-β did not, despite the fact that the amino acid sequences of their catalytic subunits are nearly identical. This finding matches our previous observation that deletion of PKA-Cat-α, but not PKA-Cat-β, resulted in loss of AQP2 expression (13). The finding was attributed to differences in the subcellular localizations of PKA-Cat-α and PKA-Cat-β, which could be due to differential interactions with PKA regulatory subunits (and associated AKAP proteins) or PDZ-domain proteins. Of the four PKA regulatory subunits (coded by Prkar1a, Prkar1b, Prkar2a and Prkar2b), only Prkar1a (PKA-RIα) was identified as a negative regulator of *Aqp2* gene transcription despite the fact that all four are expressed. This finding could be due in part to specialized localization of PKA-RIα and/or compartmentation of cAMP action (25–27). Such compartmentation could allow targeted phosphorylation of the three CREB-family transcription factors (CREB1, CREM, ATF1) that mediate regulation of *Aqp2* gene transcription (17).

### Dyrk1a

The kinome CRISPR screen preliminarily identified Dyrk1a as a strong positive regulator of vasopressin-dependent *Aqp2* transcription (**Figure 2**). To address this finding further, we used CRISPR/Cas9 genome editing to make Dyrk1a-null clonal cell lines in mpkCCD cells targeting Exon 8, which overlaps the kinase catalytic domain (**Figure 3A**). Consistent with a positive regulatory role of Dyrk1a, Dyrk1a KO cells no longer responded to vasopressin with an increase in AQP2 protein abundance (**Figure 3, B and C**). Although the cells formed a confluent monolayer, the transepithelial resistance (TER) was attenuated in Dyrk1a KO cells (**Figure 3D**). Labeling of ZO-1 (a tight junction marker) and DAPI (a nuclear marker) confirmed that the cells retain their polarized epithelial morphology but was indicative of greater local variability of cell size, measured as nuclei per unit area using the DAPI signal (**Figure 3, E and F**). FACS analysis of propidium iodide labeled cells was indicative of a partial shift from G0/G1 to the S and G2 phases, consistent with increased entry into the cell cycle (**Figure 3G**).

**Figure 3.**
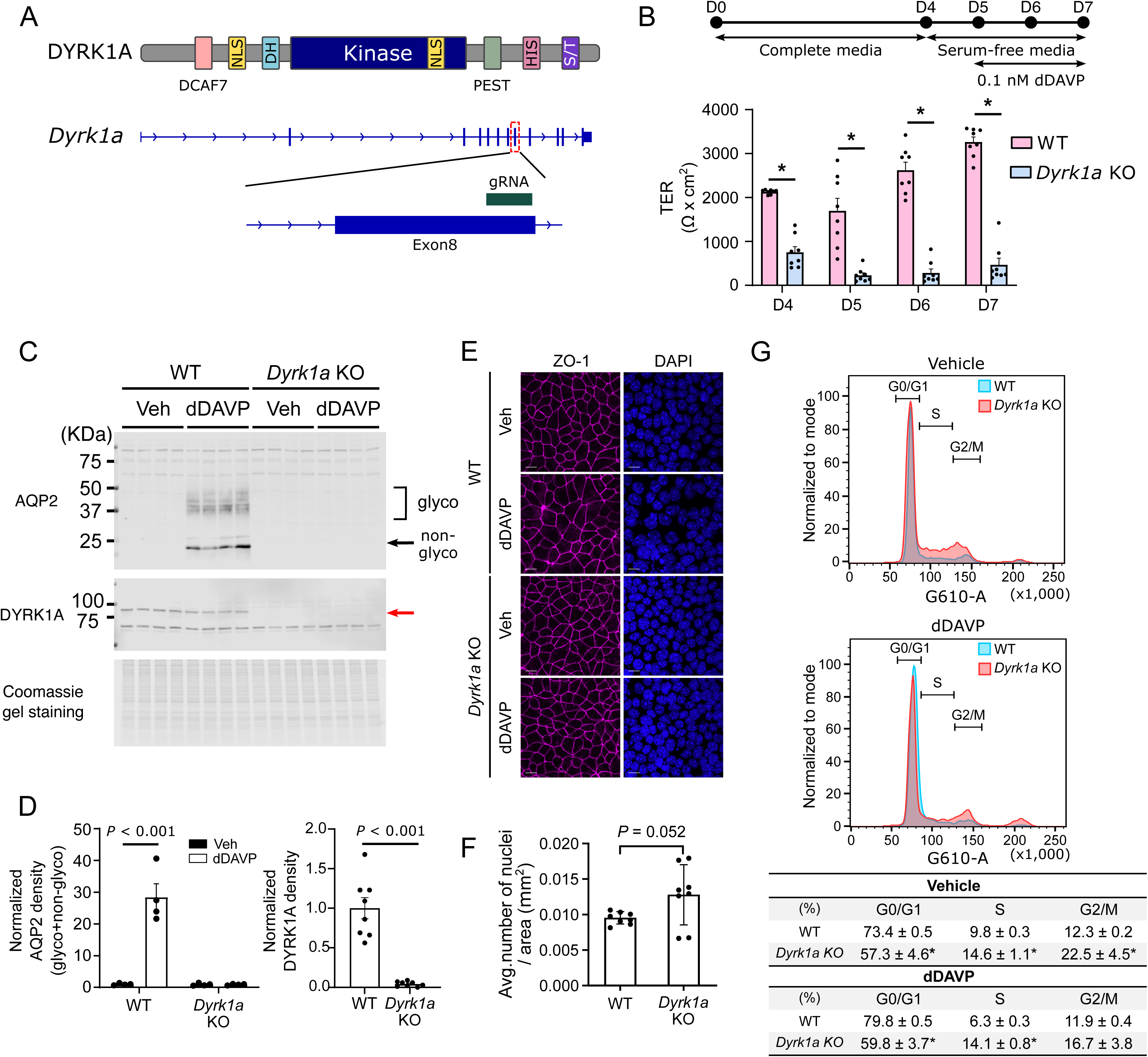
Effect of Dyrk1a deletion in clonal cell lines. **A.** DYRK1a protein and *Dyrk1a* gene structure. Protein domains: DCAF7, **<u>D</u>**DB1 and **<u>C</u>**UL4 **<u>A</u>**ssociated **<u>F</u>**actor **<u>7</u>**domain; NLS, Nuclear Localization Signal; DH, **<u>D</u>**bl **<u>H</u>**omology domain; PEST, Proline/Glutamic Acid/Serine/Threonine-rich region; HIS, HIStidine-rich region; S/T, Serine/Threonine-rich region. Protein structure and gene structure are not aligned. The sgRNA used to make clonal cell lines targets a site located in exon 8 codes for a portion of kinase catalytic domain of Dyrk1a. **B and C**. Immunoblotting and densitometry of AQP2 and Dyrk1a proteins in WT (“<u>W</u>ithout <u>T</u>ransformation”) and *Dyrk1a* KO cells (4 independent clones). glyco, glycosylated band; non-glyco, non-glycosylated band; Veh, vehicle; dDAVP, desmopressin; kDa, kilodaltons. Red arrow indicates expected molecular weight of Dyrk1a (∼84 kDa). Statistical comparisons used simple t-tests. **D**. Transepithelial resistance (TER) in WT and *Dyrk1a* KO cells. D0-D7, Day 0 through Day 7. **E**. ZO-1 immunofluorescence and DAPI fluorescence labeling of cells. ZO-1 abundance (fluorescence signal integrated across multiple fields) showed no difference between WT and Dyrk1a KO cells. **F.** Average number of nuclei per unit area. DAPI-labeled nuclei were counted from three random fields for each of four WT and four *Dyrk1a* KO lines. **G.** Cell cycle profiling by propidium iodide labeling. Distributions of cells in G0/G1, S, and G2/M phases are shown. G-610-A indicates settings for FACS (G, green laser; 610, emission filter peak transmission wavelength in nm; A, detector setting). For both vehicle– and dDAVP-treated cells, Dyrk1a KO showed a significant reduction in the G0/G1 population with a corresponding increase in S and G2/M phases compared to WT. Data are presented as mean ± SEM, * *P* < 0.05.

Dyrk1a functions in cells to activate the so-called DREAM (<u>D</u>imerization partner/<u>R</u>B-like/<u>E</u>2F/<u>A</u>nd/<u>M</u>uvB) complex by phosphorylation of one of its subunits LIN52 (33). The DREAM complex is a master transcriptional regulator that functions as a global repressor of E2F target genes, which orchestrate cell-cycle progression (49). Its main role is to limit cell cycle entry and maintain a state of differentiation (cellular quiescence, G0 phase). These prior observations coupled with our FACS-based cell-cycle analysis (**Figure 3G**) point to the hypothesis that Dyrk1a deletion results in AQP2 loss from the cells through the loss of differentiation associated with cell-cycle activation. To address this possibility, we have carried out RNA-sequencing in Dyrk1a KO cells compared with WT cells, both treated with dDAVP. The full curated data set can be viewed or downloaded at https://esbl.nhlbi.nih.gov/Databases/Dyrk1a-KO-RNA-seq/. A volcano plot of the data (**Figure 4A**) shows a greater-than 90% decrease in Aqp2 mRNA abundance in the Dyrk1a KO cells (Dyrk1a-KO/WT) = –3.56), paralleling the decrease in AQP2 protein seen previously in the same cells (**Figure 3C**). Red points show values for mRNA species corresponding to a previously curated set of differentiation markers for collecting duct principal cells (https://esbl.nhlbi.nih.gov/Signaling-Pathways/PC-Diff-Markers/) and are indicative of a predominance of markers with decreased mRNA abundances. Blue points show values for mRNA species corresponding to a previously curated set of cell-cycle genes (https://esbl.nhlbi.nih.gov/Databases/Signaling-Pathways/Cell-Cycle/) and show a predominance of transcripts with increased mRNA abundances. **Figure 4B and 4C** show the results of formalized analyses supporting the conclusion that deletion of Dyrk1a upregulates expression of cell-cycle genes while resulting in loss of differentiation markers. GO term and KEGG Pathway enrichment analyses of upregulated transcripts and heat map comparison of cell-cycle transcript abundances in individual replicates also support the conclusion that there is widespread transcriptional activation of cell-cycle genes in Dyrk1a KO cells (Supplemental Figure 1). Overall, the findings support the conclusion that Dyrk1a is a positive regulator of *Aqp2* gene expression and that a likely mechanism is a Dyrk1a-mediated shift from the cell cycle to quiescence with increased collecting duct principal cell differentiation.

**Figure 4.**
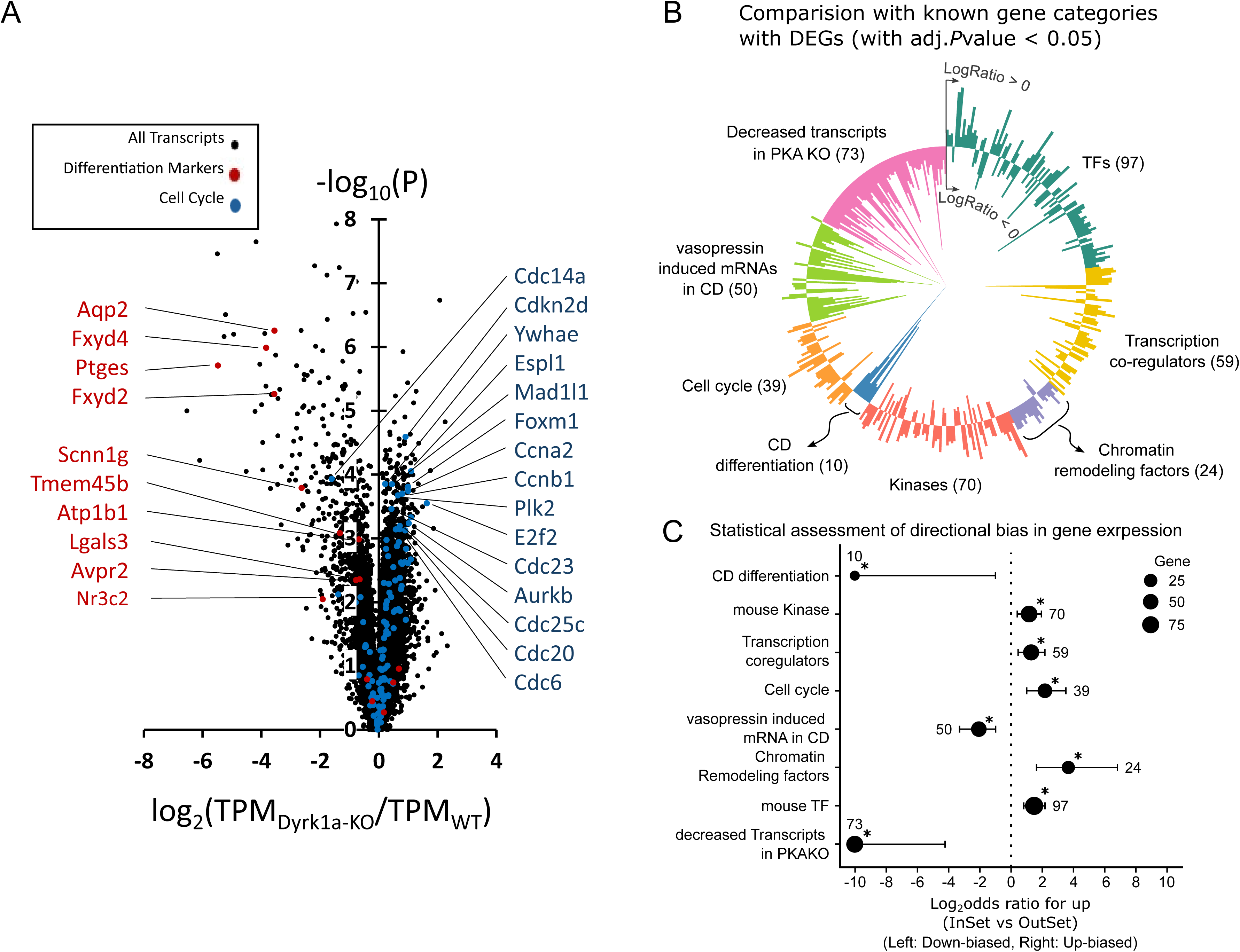
RNA-sequencing in dDAVP-treated Dyrk1a KO cells (four independent clones versus WT cells (also n=4). Cells were exposed to 0.1 nM dDAVP for 48 hrs. **A.** Volcano plot. Each point corresponds to a different mRNA species. Red points correspond to previously curated cell differentiation markers for collecting duct principal cells (https://esbl.nhlbi.nih.gov/Signaling-Pathways/PC-Diff-Markers/) (top 10 –log_10_[P] values are labeled). Blue points correspond to previously curated cell-cycle genes (https://esbl.nhlbi.nih.gov/Signaling-Pathways/Cell-Cycle/) (top 15 –log_10_[P] values are labeled). Here P is the standard P value derived from the t-statistic, used as a stratification parameter rather than a criterion of statistical significance. TPM, transcripts per million. **B.** Radial plot showing changes of differentially expressed genes (DEGs) classified into gene categories. Gene lists for categories are from https://esbl.nhlbi.nih.gov/Databases/Signaling-Pathways/ and https://esbl.nhlbi.nih.gov/Databases/KSBP2/Targets/CategorizedGeneLists.html. Bars extending outward represent upregulated transcripts, while those extending inward show downregulated transcripts. Numbers of transcripts in individual categories are indicated in parentheses. **C.** Statistical assessment of directional bias in gene expression across categories shown in B. Positive log-odds ratios indicate enrichment for upregulated transcripts, and negative values indicate enrichment for downregulated transcripts. Horizontal bars show 95% confidence intervals (CIs); dots indicate the estimated log_2_ odds ratios for each category. Log_2_OR(odds ratio) = Log_2_((Up_category_/Down_category_)/(UP_non-category_/Down_non-category_)), non-category refers to DEGs outside the tested category. An asterisk (*) denotes P < 0.05 after Benjamini–Hochberg correction. Values and CIs are displayed on a log_₂_ scale and clipped at ±10 for readability.

### Tgfbr1, Tgfbr2, and Tgfbr3

The TGF-β receptor is a well-characterized receptor kinase. Our kinome CRISPR screen preliminarily identified three TGF-β receptor subunits (Tgfbr1, Tgfbr2, and Tgfbr3) as negative regulators of vasopressin-dependent *Aqp2* transcription (**Figure 2**). Tgfbr1 and Tgfbr2 form the main receptor complex and couple with the ancillary receptor protein Tgfbr3 (which lacks independent kinase activity) (50). Prior studies implicated TGF-β in the vasopressin-escape phenomenon, a response that mitigates the degree of hyponatremia in the syndrome of inappropriate antidiuresis (SIAD) by inhibiting expression of the *Aqp2* gene in collecting duct principal cells (24).

We carried out RNA-sequencing in mpkCCD cells with exposure to TGF-β1 versus vehicle (**Figure 5**). As shown in **Figure 5A**, TGF-β1 dose-dependently decreased AQP2 protein abundance in mpkCCD cells. Expression profiling with RNA-seq was carried out in dDAVP-treated mpkCCD cells exposed to either TGF-β1 (10 ng/ml) or its vehicle (both with 0.1 nM dDAVP). An 86% decrease in Aqp2 mRNA rivaled the magnitude of decrease of AQP2 protein (**Figure 5B**) [TPM values: TGF-β+dDAVP, 182±29(SD) (n=4) versus dDAVP alone, 1297±53(SD) (n=4); P<10^−7^]. Multiple collecting-duct differentiation markers (labeled in **Figure 5B**) also showed substantial decreases in mRNA abundance. The full curated RNA-seq dataset can be browsed or downloaded at https://esbl.nhlbi.nih.gov/Databases/TGF-beta-response-RNA-seq/. The RNA-seq dataset was analyzed further to identify potential mechanisms of the decrease in *Aqp2* gene expression. A prominent feature was a >50% decrease in abundance of the Avpr2 transcript (**Figure 5B**) which codes for the vasopressin V2 receptor [TPM values: TGF-β+dDAVP, 108±6 (SD) (n=4) versus dDAVP alone, 235±6 (SD) (n=4); P<10^−6^]. This finding suggests that part of the explanation for the loss of *Aqp2* gene expression in response to TGF-β is loss of V2R-ADCY6-cAMP-PKA signaling, which is previously known to be responsible for vasopressin-mediated regulation of *Aqp2* gene transcription (12,16,17).

**Figure 5.**
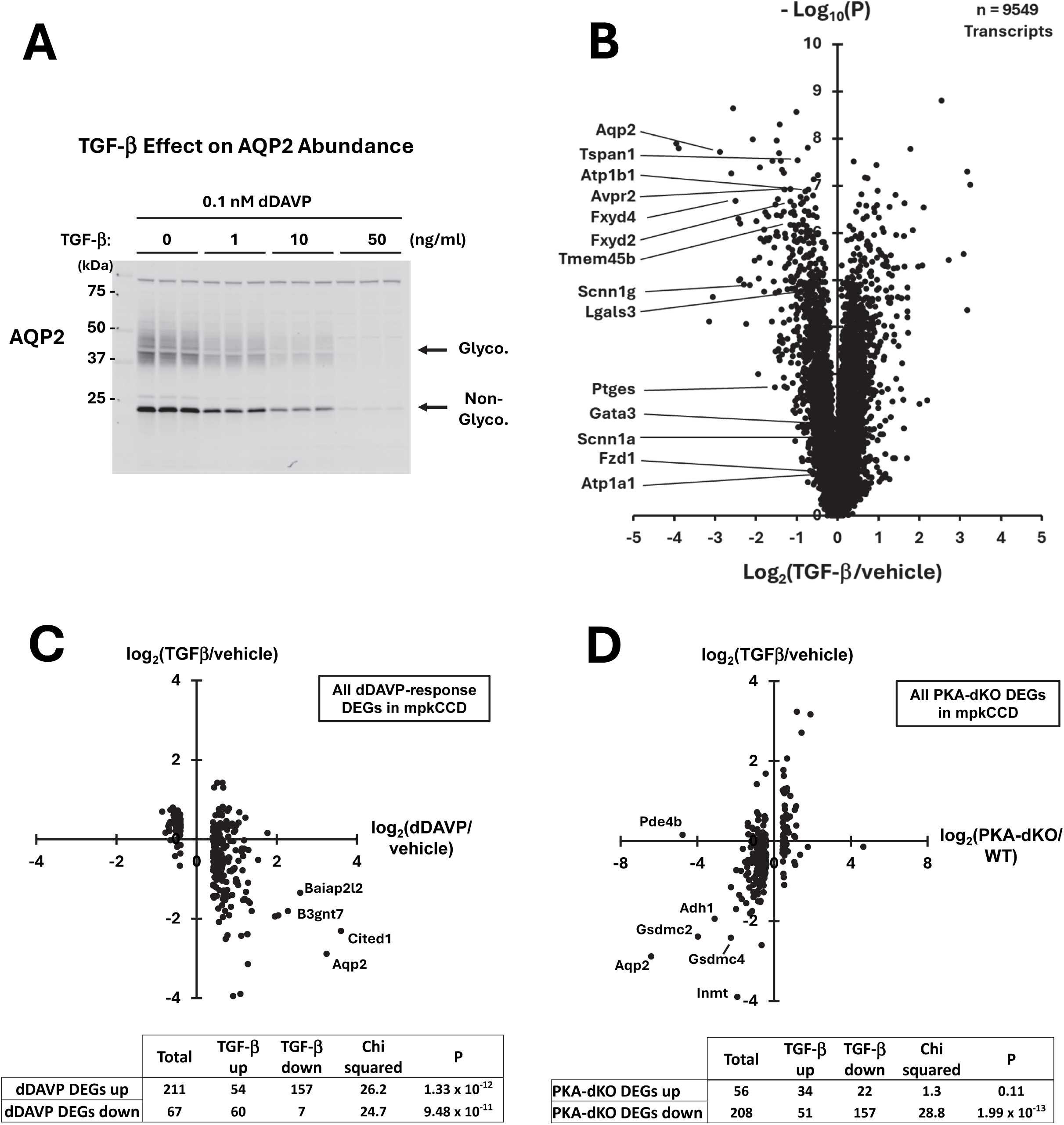
Effects of TGF-β1 in dDAVP-treated collecting duct mpkCCD cells. **A**. Western blot showing effect of TGF-β1 at indicated concentrations for 48 hours on aquaporin-2 (AQP2) protein abundance in vasopressin (dDAVP)-treated mpkCCD cells. Both non-glycosylated AQP2 (lower band) and glycosylated AQP2 (broad 35-50 kD upper band) were dose-dependently decreased. **B**. Volcano plot showing responses of 9549 mRNA species in mouse mpkCCD cells to treatment with TGF-β1 (10 ng/ml for 48 hr, n=4) versus vehicle (n=4). mRNA abundances are expressed as “transcripts per million” (TPM). All samples were maintained in 0.1 nM dDAVP in the basolateral medium. Labeled transcripts represent genes characteristic of differentiated collecting duct principal cells (https://esbl.nhlbi.nih.gov/Signaling-Pathways/PC-Diff-Markers/). **C**. Inverse relationship between dDAVP responses and TGF-β responses among all transcripts positively or negatively altered in response to dDAVP. All TGF-β response data can be viewed at https://esbl.nhlbi.nih.gov/Databases/TGF-beta-response-RNA-seq/ and the dDAVP response data can be viewed at https://esbl.nhlbi.nih.gov/Databases/Vasopressin-RNA-seq-mpkCCD/. **D**. Positive relationship between responses to deletion of both PKA catalytic subunit genes (PKA-Cat-α and PKA-Cat-β) and addition of TGF-β in WT mpkCCD cells among all transcripts significantly altered by PKA deletion. All values for TGF-β responses can be viewed at https://esbl.nhlbi.nih.gov/Databases/TGF-beta-response-RNA-seq/ and the values for effect of PKA deletion can be viewed at https://esbl.nhlbi.nih.gov/Databases/PKA-KO/Ratio.html.

If the l*oss of Aqp2* gene expression in response to TGF-β is in part due to loss of V2R-ADCY6-cAMP-PKA signaling, we would expect that additional vasopressin-induced transcripts would be regulated by TGF-β1 in a direction opposite the vasopressin response. For this, we used newly generated RNA-seq data that compares transcript abundances in dDAVP-exposed mpkCCD cells to vehicle-treated cells (https://esbl.nhlbi.nih.gov/Databases/Vasopressin-RNA-seq-mpkCCD/). Looking only at differentially expressed transcripts with dDAVP, the hypothesized inverse relationship between dDAVP responses and TGF-β responses was indeed observed (**Figure 5C**].

In addition, if the l*oss of Aqp2* gene expression in response to TGF-β is in part due to loss of V2R-ADCY6-cAMP-PKA signaling, we would expect that transcripts that change following PKA deletion (12) would exhibit parallel changes in abundance in response to TGF-β1. Considering only differentially expressed transcripts following PKA deletion, the hypothesized parallel relationship between PKA-deletion responses and TGF-β responses was indeed seen (**Figure 5D)**.

Suppression of V2R-ADCY6-cAMP-PKA signaling may not be the only cause of loss of *Aqp2* gene expression in response to TGF-β. In fact, the decrease in vasopressin V2 receptor expression requires an explanation since its expression has not been found to be regulated by its ligand, vasopressin (https://esbl.nhlbi.nih.gov/Databases/Vasopressin-RNA-seq-mpkCCD/). Because more transcripts underwent changes in abundance in response to TGF-β (n=1089) than in response to dDAVP (n=278) it appears that TGF-β affects more than just the V2R-ADCY6-cAMP-PKA pathway.

A major role of TGF-β in epithelia is mediation of epithelial-to-mesenchymal transition (EMT) (51). In EMT, epithelial-specific genes undergo decreases in expression and mesenchymal genes undergo increases, i.e. target epithelia undergo de-differentiation. The transition is often partial, reflecting the fact that the epithelial state and the mesenchymal state are endpoints on a continuum (51). The plasticity inherent in EMT is critical to tissue and organ morphogenesis including kidney development. To address the hypothesis that TGF-β induces partial EMT in mpkCCD cells, we carried out gene-set enrichment analysis to identify *Gene Ontology Biological Process* terms in which TGF-β-regulated transcripts are over-represented relative to the expected number of transcripts from random sampling from the full list of expressed mRNAs (**Figure 6, top**). If the hypothesis is true we would expect to see terms indicative of epithelial dedifferentiation, consistent with “epithelial cell differentiation regulation” and “epithelial cell proliferation” as seen. In addition, we would expect terms related to epithelial polarity, e.g. “cell junction assembly” and “regulation of cell-cell adhesion”, as seen. Importantly, the partial EMT state plays a critical role in epithelial development consistent with *Gene Ontology Biological Process* terms “tube morphogenesis”, “epithelial cell morphogenesis”, “ureteric bund development”, “canonical Wnt signaling pathway” (**Figure 6, top**). Further, a similar analysis of *Gene Ontology Cellular Component* terms (**Figure 6, middle**) identified “apical plasma membrane” and “basal plasma membrane” as the terms most highly enriched in TGF-β−regulated transcripts, consistent with the hypothesized effects on apical-basal polarity. Gene-set enrichment analysis to identify KEGG Pathways in which TGF-β−regulated transcripts are over-represented relative to the expected number of transcripts from random sampling identified “TGF-beta signaling” as expected, but also “Hippo signaling”, “Wnt signaling” and “MAPK signaling” (**Figure 6, bottom)**. The Hippo signaling pathway provides a signaling connection between cell-cell contacts at cell junctions and the transcriptional regulation of cell differentiation (52). The canonical Wnt pathway plays a key role in collecting duct development and the RNA-seq data identified three Wnt homologs as TGF-β regulated transcripts, viz. Wnt7a, Wnt9a and Wnt10a.

**Figure 6.**
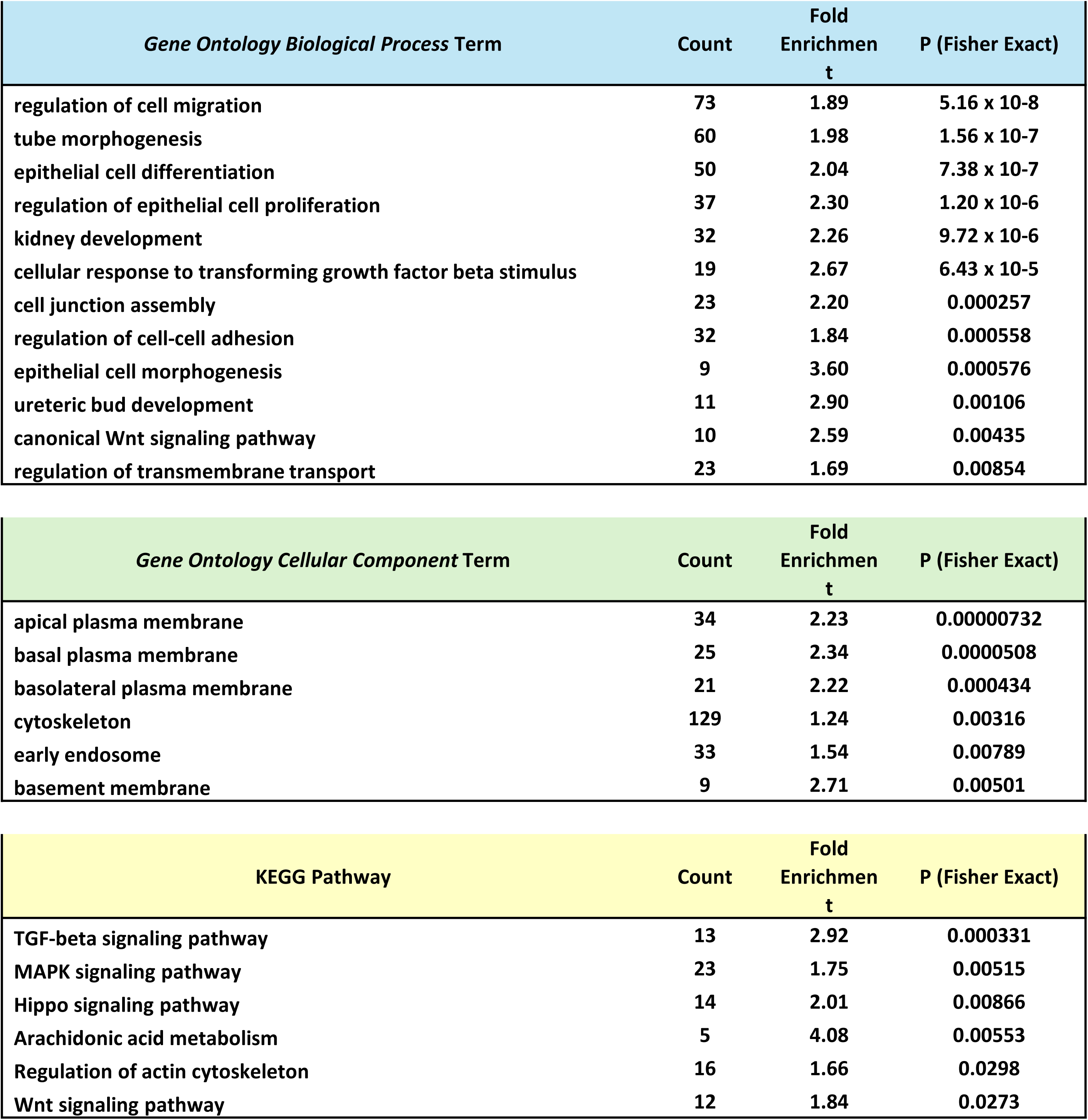
Gene-set enrichment analysis. Identification of *Gene Ontology Biological Process* terms (top), *Gene Ontology Cellular Component* terms (middle) and KEGG Pathway terms (bottom) for which TGF-β-regulated transcripts are over-represented relative to the expected number of transcripts from random sampling from the full list of expressed mRNAs.

The role of MAPK signaling is of special interest because three protein kinases in the JNK phosphorylation pathway (Map3k1, Map2k4 and Map2k7) were identified as negative regulators of *Aqp2* gene transcription in the CRISPR KO screen (**Figure 2**). Although TGF-β signals chiefly through Smad2/Smad3 phosphorylation, a “non-canonical” TGF-β signaling pathway has been shown to involve activation of the JNK pathway (53). The main phosphorylation target of JNK is the bZIP transcription factor c-Jun (54) and Jun mRNA was found to be markedly increased in abundance in response to TGF-β (TPM values: TGF-β, 39.8±3.7(SD) (n=4); vehicle, 129.0±11.2(SD) (n=4), P=5.2×10^−6^) (https://esbl.nhlbi.nih.gov/Databases/TGF-beta-response-RNA-seq/). An increase in Jun transcript abundance is an expected finding in response to increased JNK-mediated Jun activation because the Jun protein positively regulates *Jun* gene transcription in a positive feedback loop (55).

### Stk11/LKB1

The kinome CRISPR screen identified Stk11, the catalytic component of the LKB1 (Liver Kinase B1) complex, as well as an LKB1 complex accessory protein Strada, as putative strong negative regulators of vasopressin-dependent *Aqp2* transcription (**Figure 2**). To address this finding further, we used CRISPR/Cas9 to make Stk11-null clonal cell lines in mpkCCD cells, using two sgRNAs targeting the kinase catalytic domain in different clones (**Figure 7A**). The mutant cells formed confluent monolayers with high transepithelial resistance (TER), characteristic of the mature collecting duct epithelium (**Figure 7B**). Immunoblotting for aquaporin-2 (AQP2) confirmed the inferred regulatory role of Stk11 (**Figure 7C,3D**). Specifically, Stk11 deletion increased the abundance of AQP2 protein in vasopressin-treated cells. Furthermore, Stk11 deletion was associated with clear expression of AQP2 even without vasopressin stimulation, whereas AQP2 protein was not detectable in control cells in the absence of vasopressin. This finding points to LKB1 signaling as a potential target for treatment of vasopressin resistance syndrome (nephrogenic NDI).

**Figure 7.**
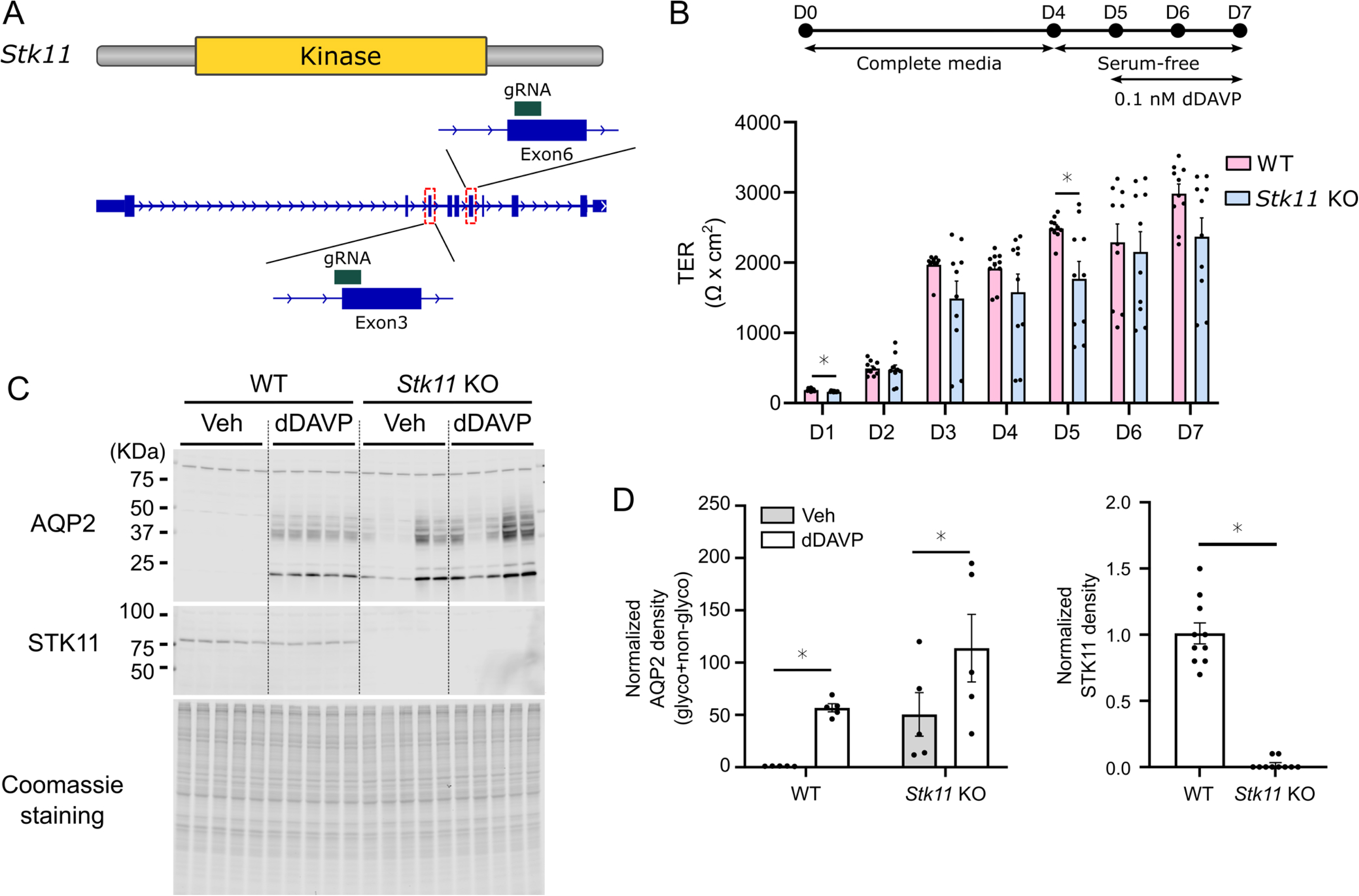
Stk11/LKB1 deletion clonal cell lines. **A**. Two sgRNAs were employed for CRSPR-Cas9 mediated targeting of DNA regions corresponding to the Stk11/LKB1 kinase catalytic domain (Exons 3 and 6) resulting in five independent Stk11/LKB1 deletion clonal cell lines for further study. **B**. Transepithelial resistance (TER) measurements in the Stk11 KO lines and in the five WT (“without transformation”) clones. D0-D7, Days 0 through 7 of culture. *, P<0.05 for simple t-test. **C.** Immunoblotting for AQP2 and Stk11/LKB1 in WT and Stk11/LKB1 deletion lines. Cells were exposed to the vasopressin analog dDAVP (0.1 nM for 48 hours) or the corresponding vehicle (Veh) as indicated. **D.** Densitometry values from immunoblotting. For AQP2, the densitometry values for the glycosylated (37-28 kDa) and non-glycosylated bands (∼24 kDa)were summed. *, P<0.05 for simple t-test.

To test whether the Stk11 deletion also increases the abundance of Aqp2 mRNA and other known vasopressin-regulated genes, we carried out RNA-seq in Stk11 KO (n=5) versus WT (n=5) cells in the absence of vasopressin. The full curated RNA-seq data can be viewed at https://esbl.nhlbi.nih.gov/Databases/Stk1-KO-no-VP/. Indeed, in the absence of vasopressin, there was a marked increase in Aqp2 mRNA abundance (TPM values: WT, 190±12; Stk11 KO, 5126±1751, FDR-corrected P value, 0.000004), confirming the vasopressin-like response of Stk11 deletion seen by immunoblotting. **Figure 8A** shows a plot of all data for the Stk11 knockout vs WT cells in the absence of vasopressin and **Figure 8B** shows RNA-seq data for a parallel experiment to assess the response to the vasopressin analog dDAVP vs vehicle in WT cells (Full curated data for dDAVP vs vehicle at https://esbl.nhlbi.nih.gov/Databases/Vasopressin-RNA-seq-mpkCCD/). The red points in **Figure 8A** indicate values for the 60 transcripts found to be increased in response to dDAVP ([Log_2_(TPM_dDAVP_/TPM_Vehicle_] to values greater than 0.8 **Figure 8B**). **Figure 8C** shows changes in individual mRNA species in response to Stk11 deletion (vertical axis) plotted against changes seen in response to exposure of WT cells to dDAVP (horizontal axis), illustrating the similarity of the two responses. Overall, these data indicate that Stk11 deletion mimics the effect of activation of the V2R-G_α_s-ADCY6-cAMP-PKA signaling pathway.

**Figure 8.**
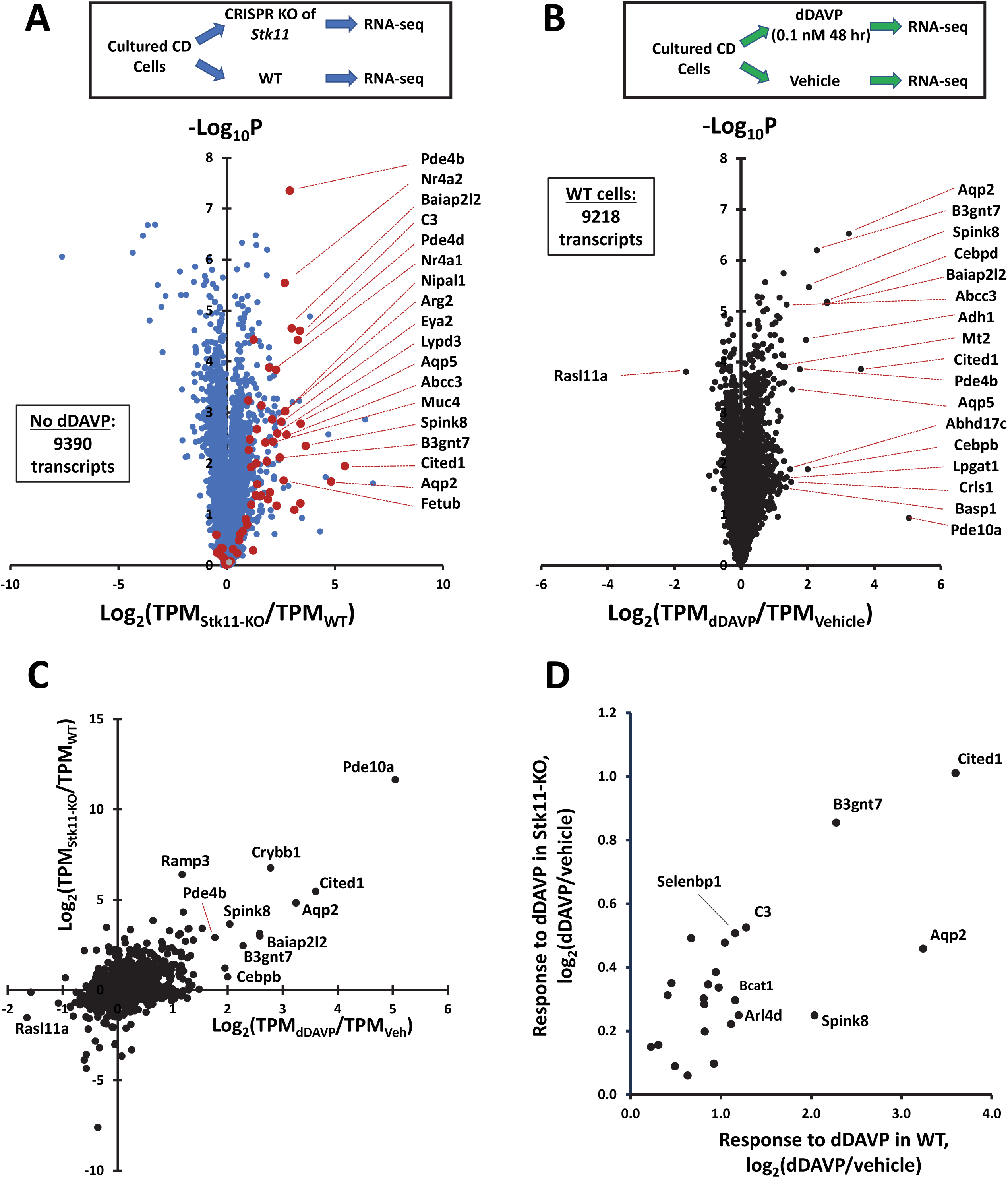
Response to deletion of Stk11 (LKB1) mimics the response to vasopressin analog dDAVP. **A.** RNA-seq data comparing cultured collecting duct cell (mpkCCD) clones after CRISPR knockout of Stk11 (Stk11-KO) versus control clones without Stk11 deletion (WT, ‘without transformation’). Cells were not exposed to dDAVP. Horizontal axis shows base 2 logarithm (“Log_2_“) of ratio TPM_Stk11-KO_/TPM_WT_ for individual mRNA species. TPM, transcripts per million. Vertical axis shows the negative of the base 10 logarithm of the P parameter derived from the t-statistic for n=5 versus n=5 comparison for each mRNA species. Data were filtered to include only the 9390 most abundant transcripts (mean TPM for either Stk11-KO or WT >10). Transcripts that are increased in abundance in mpkCCD cells in response to dDAVP (from data in Figure 4B) are indicated as red points. Official gene symbols for transcripts with the highest values of Log_2_(TPM_Stk11-KO_/TPM_WT_) are labeled at right. Full data set available at https://esbl.nhlbi.nih.gov/Databases/Stk1-KO-no-VP/. **B.** RNA-seq data comparing cultured collecting duct cells (mpkCCD) exposed to dDAVP (0.1 nM) for 48 hours with vehicle control values for the same clonal mpkCCD lines. Horizontal axis shows base 2 logarithm (“Log_2_“) of ratio TPM_dDAVP_/TPM_Vehicle_ for individual mRNA species. Vertical axis shows the negative of the base 10 logarithm of the P parameter derived from the t-statistic for n=5 versus n=5 comparison for each mRNA species. Data were filtered to include only the 9218 most abundant transcripts (mean TPM for either dDAVP or vehicle >10). Selected official gene symbols for responding transcripts are shown at right. Complete dataset available at https://esbl.nhlbi.nih.gov/Databases/Stk11-KO-RNA-seq/. These data are highly concordant with data from Sandoval et al (16). **C.** Comparison of responses to Stk11 deletion and dDAVP administration in mpkCCD cells. Linear regression: y = 0.957x + 0.022 where y is Log_2_(TPM_Stk11-KO_/TPM_WT_) and x is Log_2_(TPM_dDAVP_/TPM_Veh_). P < 10^−8^ (slope value versus zero) for all 9162 transcripts with values in both data sets. Selected transcripts are labeled with corresponding gene symbols. **D.** Effect of Stk11 deletion on response to dDAVP. Data presented are for the 24 transcripts reported to be increased in abundance in response to dDAVP by Sandoval et al (16). dDAVP-mediated increases were smaller in Stk11 KO cells than in WT cells for all 24 transcripts. Full data can be viewed in **Supplemental Table 1** including P values for dDAVP responses for each of the 24 transcripts.

The foregoing observations establish that Stk11/LKB1 and PKA have opposing effects on *Aqp2* gene expression. To explore the nature of the interaction between Stk11/LKB1 and PKA, we asked if Stk11 expression is necessary for V2R-G_α_s-ADCY6-cAMP-PKA-mediated responses. To address this, we use RNA-seq data to compare the vasopressin response in Stk11/LKB1 KO lines versus WT controls. The full curated data sets are provided at https://esbl.nhlbi.nih.gov/Databases/Stk11-KO-RNA-seq/. **Figure 8D** summarizes data for 24 mRNA species previously found to be significantly increased in abundance by vasopressin (16). All of these transcripts showed significant increases in WT cells in response to the vasopressin analog (log_2_[dDAVP/vehicle] > 0 and P < 0.05). The responses in the Stk11/LKB1 KO cells were markedly attenuated but not eliminated for most transcripts. All mRNA species had smaller log_2_[dDAVP/vehicle] values in the Stk11/LKB1 KO cells. (See **Supplementary Table 1** for detailed statistical analysis.) This finding supports the role of Stk11/LKB1 as an integral determinant of vasopressin V2 receptor signaling. Even though the vasopressin responses were attenuated in the Stk11-KO cells, the fact that significant responses were seen suggests that Stk11 signaling is not required for some element of the vasopressin response.

The existing literature provides a plausible hypothesis about how Stk11/LKB1 inhibits *Aqp2* gene expression (**Figure 9A**). We know that *Aqp2* gene transcription is dependent on CREB-family transcription factors (CREB1, ATF1 and CREM) (17). The activities of each of the three CREB-family transcription factors (represented by CREB1 in **Figure 9A**) are known to be regulated in two ways, both dependent on PKA: 1) Direct PKA-mediated phosphorylation of a key serine in the so-called KID domain (Ser133 in CREB1) (56) that facilitates binding of a transcriptional co-factor CREB-binding protein (CBP) (57); and 2) variable activation by one or more members of the cAMP-responsive transcriptional coactivator (CRTC) family (also known as TORCs, or Transducers of Regulated CREB activity) (58). CRTC family proteins are negatively regulated by salt-inducible kinases (SIKs) through 14-3-3-mediated sequestration in the cytoplasm (59). The SIKs are members of the SNF1 family of protein kinases that are activated by Stk11/LKB1-mediated active site phosphorylation (60). PKA-mediated phosphorylation negatively regulates SIKs resulting in net activation of CRTCs through enhancement of binding to the bZIP domains of the three CREB-family transcription factors (**Figure 9A**) (59,61).

**Figure 9.**
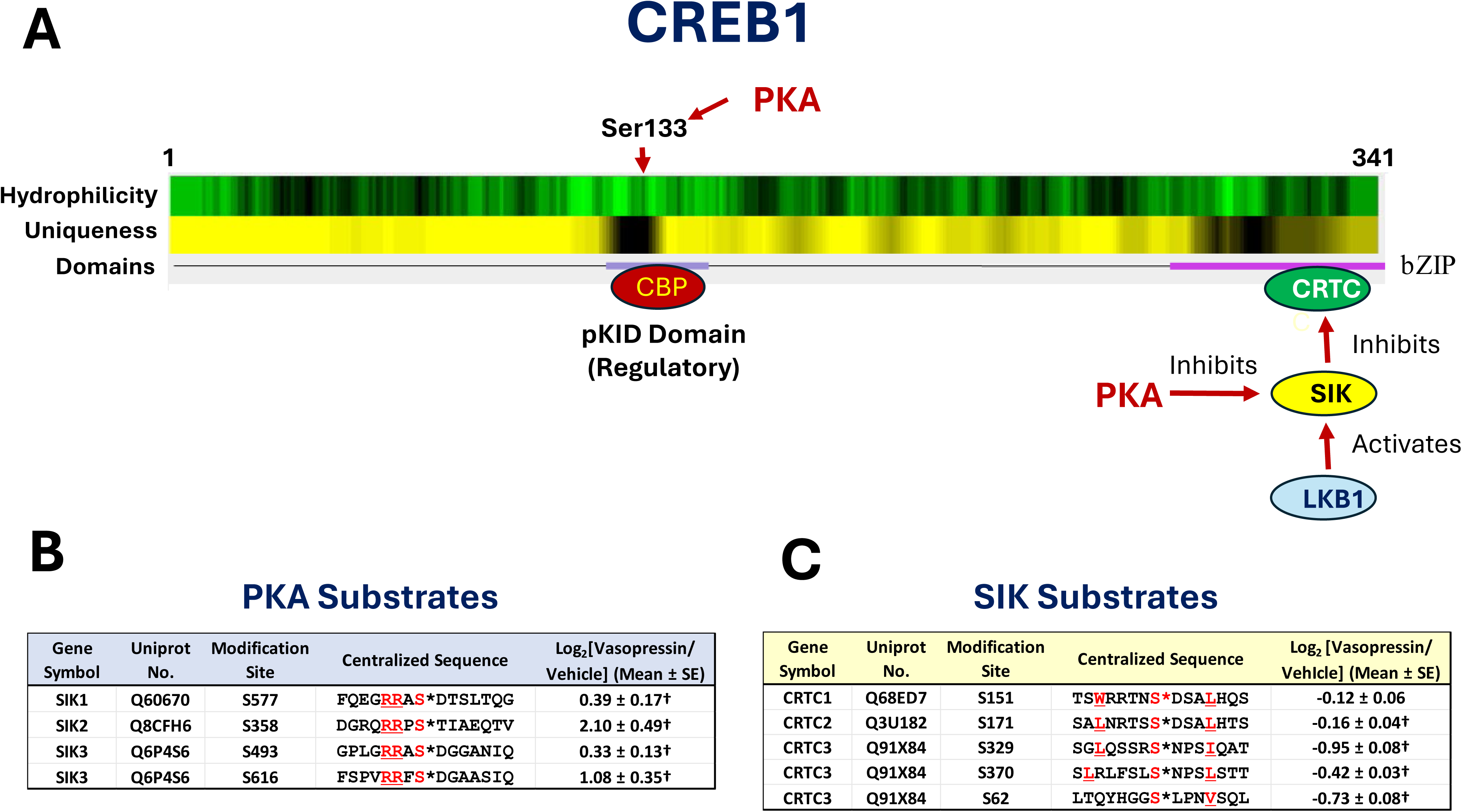
Regulation of CREB-family transcription factors by PKA and LKB1. **A**. Model of CREB1 activity regulation by PKA based on prior literature. See text for explanation and abbreviations. The other two members of the CREB family (ATF1 and CREM) are likely regulated in the same manner. Hydrophilicity (Kyte-Doolittle) and Sequence Uniqueness were mapped using *AbDesigner* (https://esbl.nhlbi.nih.gov/AbDesigner/). **B.** Phosphoproteomic data showing effect of short-term vasopressin exposure on phosphorylation of salt-inducible kinases (SIKs) at known PKA target sites in mpkCCD cells. **C.** Phosphoproteomic data showing effect of short-term vasopressin exposure on phosphorylation of CRTC proteins at known SIK target sites in mpkCCD cells. In B and C, values are from Datta et al. (22) representing 3 SILAC pairs, downloaded from https://esbl.nhlbi.nih.gov/Databases/mpkCCD-AVP/. * indicates the phosphorylation site in the centralized sequences listed. † signifies that in the original paper, the indicated log_2_(Vasopressin/Vehicle) value was considered significantly different from 0.0 with P<0.05.

The applicability of the model in **Figure 9A** to V2R-G_α_s-ADCY6-cAMP-PKA signaling in collecting duct cells and regulation of *Aqp2* gene transcription is supported in part by prior phospho-proteomic results (**Figure 9 B and C**). To address whether vasopressin signaling is associated with increased phosphorylation in the three SIK proteins, we mined data from a prior phospho-proteomic study that used quantitative protein mass spectrometry to identify phosphorylation events triggered by exposure of mpkCCD cells to the vasopressin analog dDAVP (**Figure 9B**) (22). Indeed, known PKA-target sites (R-R-X-pS motif) in SIK1, SIK2, and SIK3 proteins showed significant increases in phosphorylation in response to vasopressin. Similarly we mined data from the same study quantifying changes in phosphorylation at known SIK target sites in the three CRTC proteins (**Figure 9C**) (22). Indeed, known SIK target sites in CRTC2 and CRTC3 (59,62–65) underwent decreased phosphorylation in response to vasopressin with the largest effects on CRTC3 phosphorylation.

The model shown in **Figure 9A** provides a framework that can potentially explain the key findings of the present study with regard to Stk11/LKB1 and V2R-Gαs-ADCY6-cAMP-PKA regulation of *Aqp2* gene expression. First, the finding that Stk11 deletion mimics the action of vasopressin to increase *Aqp2* gene expression follows directly from the model. The model provides a mechanistic basis for thinking about potential therapeutic strategies for vasopressin-resistance syndromes including both genetic and acquired nephrogenic diabetes insipidus. Second, the existence of two PKA-regulated pathways (**Figure 9A**), only one of which is dependent on a priming action of Stk11/LKB1, can explain why Stk11 deletion attenuates but does not entirely eliminate the response to vasopressin.

### Overall summary

This study confirms the tremendous power of a CRISPR/Cas9 KO screening approach to identify heretofore unknown components of signaling pathways. However, it is important to recognize some important limitations of the approach. First, CRISPR screening can produce false positive results due to off-target effects, but this is mitigated by use of multiple sgRNAs per target. Our kinase library has 4 independent sgRNAs per target. Second, there is potential for false-negative results in cases where redundancy exists. The technique depends on use of a high degree of dilution of sgRNA-bearing viral particles so that no cell is transformed simultaneously by more than one sgRNA. Thus, in the present study, if the kinome sgRNA library targets two protein kinases with equivalent function, a false negative could be reported out for both. Interestingly, this did not happen with the two PKA catalytic subunit genes (*Prkaca* and *Prkacb*), confirming our prior conclusion that despite nearly identical catalytic regions, the two corresponding proteins PKA-Cat-α and PKA-Cat-β have distinct regulatory targets (13). Overall, we report 26 kinase ‘hits’ that are associated with either positive or negative effects on *Aqp2* gene transcription (**Table 1**). We validated only a subset of these in focused studies (**Figure 2C**), leaving the majority for future work. Based on the work presented here, it appears the effect of Dyrk1 deletion on *Aqp2* gene expression is not due to a specific interference with V2R-G_α_s-ADCY6-cAMP-PKA signaling, but rather is a result of a broad effect on differentiation. In contrast, both the TGF-β receptor kinase and LKB1 (Stk11 and it accessory protein Strada) interact directly with different aspects of V2R-G_α_s-ADCY6-cAMP-PKA signaling to inhibit *Aqp2* gene expression. Besides the translational possibilities of these findings for management of water balance disorders, an important long term goal is to address the generalizability of the findings to other G_α_s-coupled GPCRs.

## Methods

### Cell line and cell culture

Mouse cortical collecting duct cell line (mpkCCDc11 secondary clone 38, mpkCCD_c11-38_) has been utilized in this study. Cells were cultured in DMEM/Ham’s F12 mediaum (DMEM/F12, Gibco) containing 5 μg/mL insulin, 50 nM dexamethasone, 1 nM triiodothyronine, 10 ng/mL epidermal growth factor, 60 nM sodium selenite, 5 μg/mL transferrin, and 2% fetal bovine serum which is referred to complete media at 37°C. To polarize the cells, cells were maintained on permeable membrane filter (3450 and 3460, Corning) in complete medium for 4 days. Then, cells were incubated in serum-free/hormone-deprived medium [containing 50 nM dexamethasone, 60 nM sodium selenite, 5 μg/mL transferrin] which is referred to simple medium for another 24 hours before 1-desamino-8-D-arginine-vasopressin (dDAVP)[V1005, Sigma] treatment. To stimulate AQP2 transcription, cells were treated with dDAVP at the basolateral side of polarized cells for 48 hours in simple medium.

### Cas9-expressing GFP-AQP2 reporter cell line

GFP-AQP2 reporter mpkCCD cells were generated as previously described (23). To generate the Cas9-expressing GFP-AQP2 reporter line, Cas9 lentivirus was produced from Addgene plasmid (#52962) and transduced to GFP-Aqp2 reporter cells (Clone 43) to generate Cas9-expressing GFP-Aqp2 cells (mpkCCD_43-16A_ cells).

### Amplification of kinome library and lentivirus production

Mouse kinome wide CRIPSR knockout pooled library was obtained from Addgene (Mouse Kinome CRISPR Knockout Library (Brie), Cat. 75316, Addgene) which contains 2852 unique gRNAs targeting 713 mouse kinase genes with four gRNAs per target (PMID 26780180). Library was amplified using One Shot™ Stbl3™ Chemically Competent E. coli (C737303, Invitrogen). The transformation efficiency was calculated, and the library was confirmed that it maintained sufficient coverage (> 500 colonies per gRNA). The library was extracted using ZymoPure II Plasmid Midiperp Kit (4201, Zymo Research) according to the manufacturer’s instruction. To generate viral particles containing each gRNAs, lenti-X HEK293T cells were transfected with C3 psPAX2 (packaging plasmid), C1 pMD.2G (envelope plasmid), and kinome library plasmid. Media containing viral particles were collected twice, at 48 and 72 hours post-transfection. The media were centrifuged at 1,000 rpm for 5 minutes to remove debris, then passed through a 0.45 µm syringe filter, and stored at –80°C.

### Transduction of gRNA containing lentiviral particle and Fluorescence-activated cell sorting

Two rounds of viral tittering across dilution ratios, and a 1:320 dilution yielded an MOI of ∼0.1. For library transduction, mpkCCD_43-16A_ cells were seeded at T255 flasks at 1 x 10^6^ cells per flask (1.5 × 10^7^ cells total) and exposed to virus-containing medium at an multiplicity of infection (MOI) of 0.1, providing ∼500 transduced cells per guide with polybrene (8mg/ml) and blasticidin (8 ug/ml). Transduced cells were selected with puromycin (2 μg/ml) for 6 days. Then, cells we seeded at 6-transwell filter system and maintained in complete media for 4 days, followed by 24 hours starvation period in simple medium. Subsequently, cells were treated with 1 nM dDAVP for 48 hours in simple medium. Cells were trypsinized and sorted into four groups based on GFP intensity: GFP-negative, GFP-low, GFP-medium, and GFP-high. Cells from each group were grown at T255 in the presence of puromicyn (1 μg/ml) until cells were confluent to retrieve 10 ∼20 million cells for genomic DNA extraction. The screen was performed twice in independent rounds under identical conditions.

### Genomic DNA extraction and DNA sequencing

Genomic DNA were extracted using Quick-DNA Midiprep Plus kit (D4075, Zymo Research) according to the manufacturer’s instruction. Genomic DNA were amplified with mixture of P5 primers with stagger regions of eight different length and nine different P7 primer with barcode sequence. PCR amplification was performed using NEBNext^®^ Ultra^TM^ II Q5^®^ Master Mix (M0544S, New England Biolabs) according to the manufacturer’s instructions. Amplicons of the expected size (453 bp) were verified by agarose gel electrophoresis and purified using AMPure XP beads (A63880, Beckman Coulter). Purified amplicons were then subjected to Single-read sequencing on Illumina NovaSeq 6000. Sequencing data were processed on Galaxy Europe (https://usegalaxy.eu/) using MAGeCK suite. Reads were demultiplexed, quality-checked (FastQC), and adapter/constant regions were trimmed (Cutadapt). Guide counts were generated with MAGeCK count against the reference sgRNA library. Normalized sgRNA counts from Galaxy-MAGeCK were further analyzed to identify differentially represented genes. Three contrasts were assessed: GFP-high vs GFP-negative, GFP-high vs GFP-low, and GFP-high vs GFP-negative/low, and each gene were summarized direction (enrichment and depletion) and significance at FDR < 0.05.

### Generation of CRISPR/Cas9 mediated Dyrk7a and Stk77 knock-out cell line

The sequences of the four gRNAs were extracted from the kinome CRISPR knockout library. The sequence of the four gRNAs are 1) *Dyrk1a* targeted gRNAs: 5’-TCAGCAACCTCTAACTAACC-3’, 5’-TCTTGATTGCACTCCGTTTG-3’, 5’-TCATTGGCACCACTGAACAG-3’, 5’-TAATAGGAGTACAAACCACC-3’, 2) *Stk11* targeted gRNAs: 5’-TCATCGGCAAGTACCTGATG-3’, 5’-ACTCCATCACCATATACGTA-3’, 5’-AGTCCATTGGCAATCTCAGG-3’, 5’-GGGCCTGTACCCATTTGAGG-3’. Each gRNA containing CMV-eCas9-2a-tGFP plasmid were generated by Sigma-Aldrich. mpkCCD_c11-38_ cells were transfected plasmid (2.5 µg) using lipofectamine 3000 (L3000015, Invitrogen) according to the manufacturer’s instructions. After 48 hours of transfection, GFP-positive cells were sorted into 96-well plates (∼1 cell per well) using a BD FACSAria cell sorter. Each clone was cultured over 2-3 weeks, and the expression of Dyrk1a and Stk11 protein was evaluated by semiquantitative immunoblotting. The indel mutation was examined by *in vitro* CRISPR/Cas9 reaction using Screen It^TM^ CRISPR Cas9 Cleavage Detection kit (Cat. G990, Abm) according to the manufacturer’s instruction. Clones which showed indel mutation in all alleles were used for the further experiments.

### Measurement of transepithelial electrical resistance (TER)

Wild-type and Dyrk1a KO cells were cultured on permeable membrane filter. The medium was changed daily, and TEER was measured before changing media from fourth day to seventh day, following the manufacturer’s instruction. Briefly, an electrode pair was immersed in both apical and basolateral side, and resistance (Ω) was obtained. The resistance was measured from four replicates of wild-type and Dyrk1a KO clones. Final resistance were calculated by subtracting blank resistance from the raw resistance.

### Semiquantitative immunoblotting and antibodies

Semiquantitative immunoblot analysis was performed as previously described (ref). Briefly, cells were rapidly washed with ice-cold Dulbecco’s phosphate-buffered saline (DPBS) and lysed with Laemmli buffer (10% SDS, 10 mM Tris-HCl, pH 6.8) containing protease/phosphatase inhibitor (Halt Protease and Phosphatase Inhibitor Cocktail 100X, 78441, Thermo Fisher Scientific), then the collected lysates were placed in QIAshredder column (79656, QIAGEN) and centrifuged at 12,000 g for 2 min. The protein concentration was measured (23227, Pierce BCA protein assay Kit, Thermo Scientific), then samples were denatured by incubating at 65°C for 15 min. The protein samples were separated by SDS-PAGE using 4-20% or 12% Criterion TGX Gels (5671095, 5671045, Bio-Rad). The proteins were transferred to nitrocellulose membranes, blocked and probed with primary antibodies. Primary antibodies were anti-AQP2 (K5007, 1:5000), anti-Dyrk1a (1:1000, CST), and Stk11 (1:1000, CST). Blocking buffer (927-70001, Li-COR) and infrared fluorescence-conjugated secondary antibodies (926-68071, Li-COR) were obtained from Li-COR. Blots were visualized by the Li-COR Odyssey System (ODY-0428) and intensities of blots were analyzed using Li-COR Image Studio software.

### Immunofluorescence Microscopy

Cells were grown on a permeable membrane filter system as described above (Cell culture section). Briefly, after 48 hours of dDAVP treatment, cells were fixed with 4% paraformaldehyde for 10 min at room temperature (RT). Cells were permeabilized with permeabilizing solution (0.3% Triton X-100 and 0.1% bovine serum albumin in PBS) for 10 min and washed three times, followed by blocking with blocking solution (1% bovine serum albumin and 0.2% gelatin in PBS) for 30 min. ZO-1 were labeled with Anti-ZO-1 antibody (1:100, 339194, Invitrogen) and nuclei were labeled with 4’,6-diamidino-2-phenylindole (DAPI). Confocal fluorescence images were obtained on Zeiss LSM 880 microscope (National Heart, Lung and Blood Institute, Light Microscopy Core Facility). For each of four WT and four *Dyrk1a* KO clones, three random fields were acquired. DAPI-stained nuclei were counted in ImageJ, and counts were normalized to the cross-sectional area (nuclei per unit area).

### RNA isolation and sequencing

WT and KO cells were grown on a permeable membrane filter system as described above (Cell culture section). Total RNA was extracted using Direct-zol RNA Microprep kit (R2062, Zymo Research) following the manufacturer’s instructions. First-strand and double-stranded cDNA was synthesized and amplified from 1 ng of total RNA using SMART-Seq® mRNA HT Kit (634796, TAKARA). Amplified cDNA were purified with the AMPure XP beads (A63880, Beckman Coulter), and quantified using the Qubit dsDNA HS DNA assay kit (Q32851, Invitrogen). One nanogram of the synthesized cDNAs were fragmented and tagged with index primers (FC-131-1024, Nextera XT DNA library Preparation Kit, Illumina) following the manufacturer’s protocol. RNA-seq was performed by 50-bp paired-end sequencing on an Illumina NovaX platform. Raw sequencing reads were aligned with STAR 2.7.10 b to the mouse reference genome (Ensembl genome 113). Transcript per million and expected counts were generated by RSEM 1.3.1. Expected counts from in WT and KO samples were analyzed for differential gene expression analysis using edgeR 3.38.4.

### Cell cycle analysis using propidium iodide (PI) staining

Cells were trypsinized and washed twice with DPBS. Equal cell numbers were used across samples (2.0 x 10^6^ cells), resuspended in 100 μl cold DPBS, and fixed by adding 1 ml of 77% ethanol dropwise while vortexing (final 70% ethanol). Following a 2 hrs incubation at 4°C, cells were centrifuged at 1,200 x g for 5 min at 4°C, and washed twice with cold DPBS. Cells were suspended in 750 μl of DPBS containing 50 μg/ml PI (P3566, Invitrogen) and 50 μg/ml RNase A (EN0531, Thermo Sientific) for overnight incubation in dark. Cells were passed through a cell strainer and analyzed by flow cytometry.

### Bioinformatics and statistics

Most analyses were carried out using Microsoft Excel (https://www.microsoft.com/en-us/microsoft-365/excel) and R software (https://www.r-project.org/).

## ACKNOWLEDGMENTS

The study utilized the NHLBI Flow Cytometry Core Facility (Pradeep Dagur, Director), the NHLBI DNA Sequence Core Facility (Yuesheng Li, Director), and the NHLBI Light Microscopy Core Facility (Christian Combs, Director).

## SUPPORTING INFORMATION

[Supplementary Table 1](https://esbl.nhlbi.nih.gov/Databases/Kinome-Screen-Supplementary/)

[Supplementary Figure 1](https://esbl.nhlbi.nih.gov/Databases/Kinome-Screen-Supplementary/)

## DATA SHARING

Curated RNA-seq datasets can be accessed at the following sites:

1. https://esbl.nhlbi.nih.gov/Databases/Kinome-CRISPR-screen/
2. https://esbl.nhlbi.nih.gov/Databases/Dyrk1a-KO-RNA-seq/
3. https://esbl.nhlbi.nih.gov/Databases/TGF-beta-response-RNA-seq/
4. https://esbl.nhlbi.nih.gov/Databases/Vasopressin-RNA-seq-mpkCCD/
5. https://esbl.nhlbi.nih.gov/Databases/Stk1-KO-no-VP/
6. https://esbl.nhlbi.nih.gov/Databases/Stk11-KO-RNA-seq/
7. https://euijungpark.shinyapps.io/kinase_screening/

Raw RNA-seq data are archived at https://www.ncbi.nlm.nih.gov/geo/query/acc.cgi?acc=GSE326336

## Figure Legends

**Supplementary Table 1.**
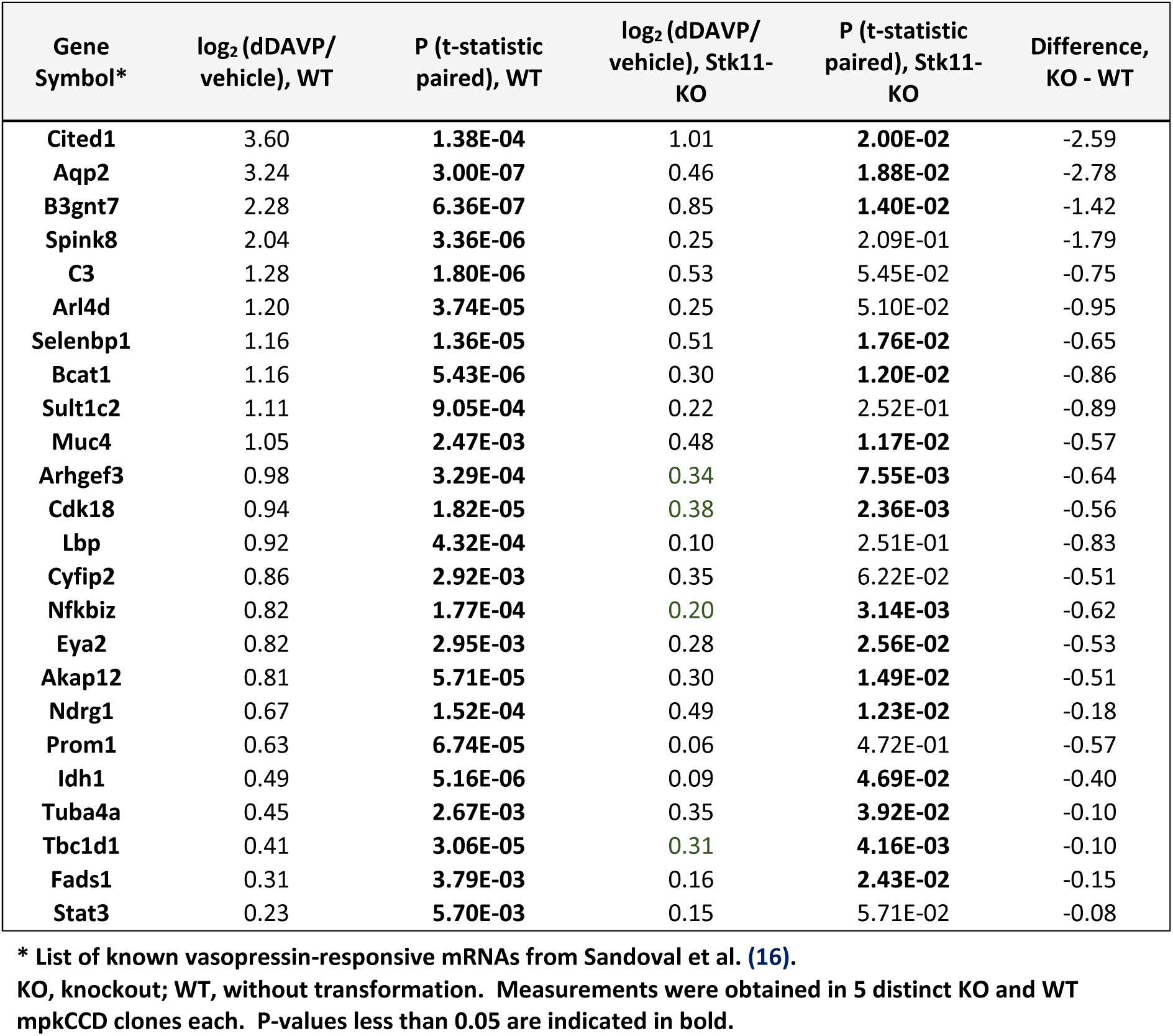
Effect of Stk11 (LKB1) knockout on response to vasopressin analog dDAVP for previously recognized vasopressin-responsive transcripts (RNA-Seq) *.

**Supplemental Figure 1.**
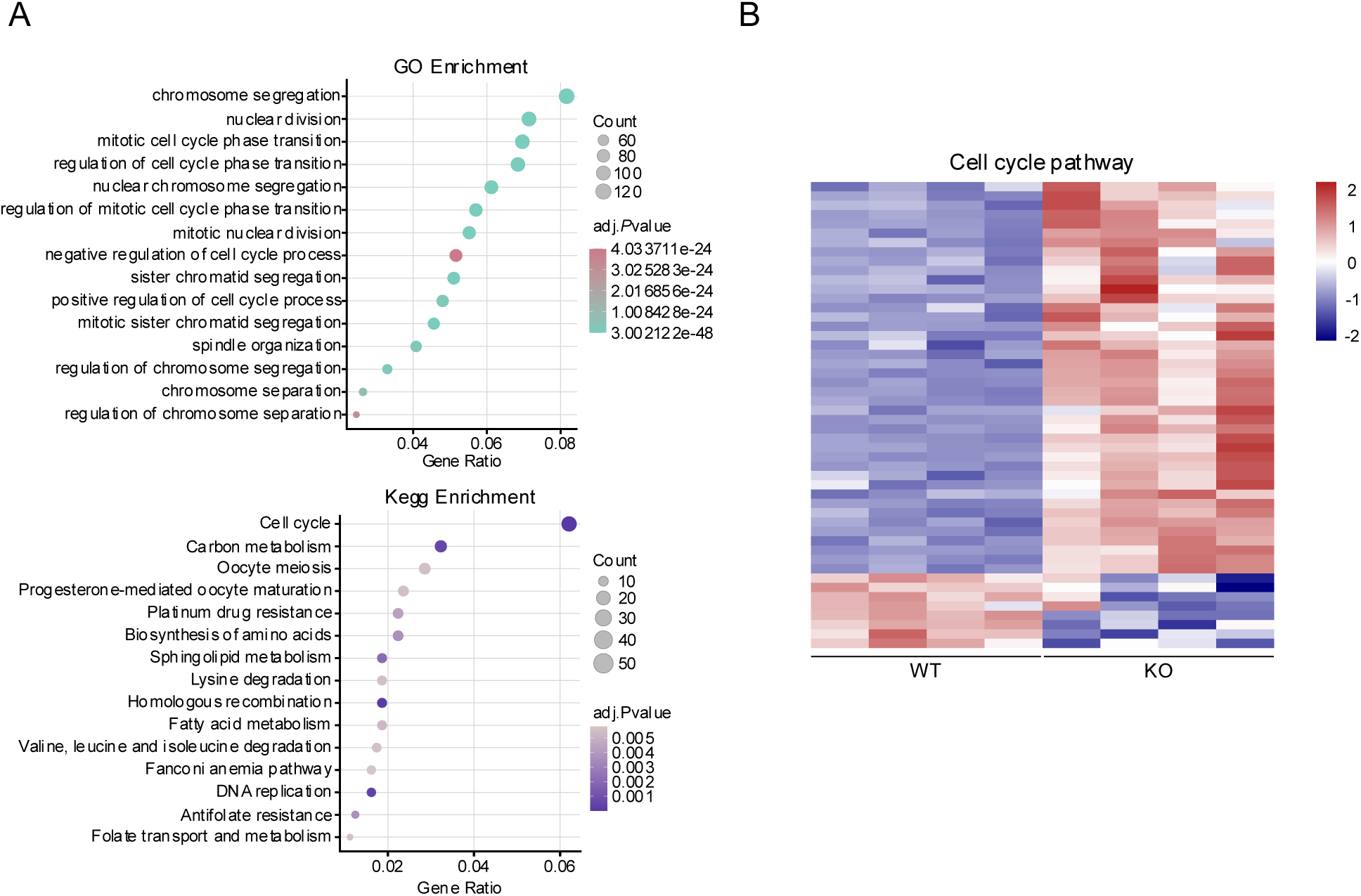
Analysis of data from RNA-sequencing of dDAVP-treated Dyrk1a KO cells (four independent clones versus WT cells (also n=4). Cells were exposed to 0.1 nM dDAVP for 48 hrs. **A.** Top 15 Gene Ontology (GO) terms and KEGG pathways from enrichment analyses of DEGs. Gene Ratio indicates the fraction of input DEGs annotated to the term (Count / total DEGs analyzed). **B.** Heatmap of cell-cycle transcripts from the KEGG Cell cycle pathway showing sample-to-sample variability.

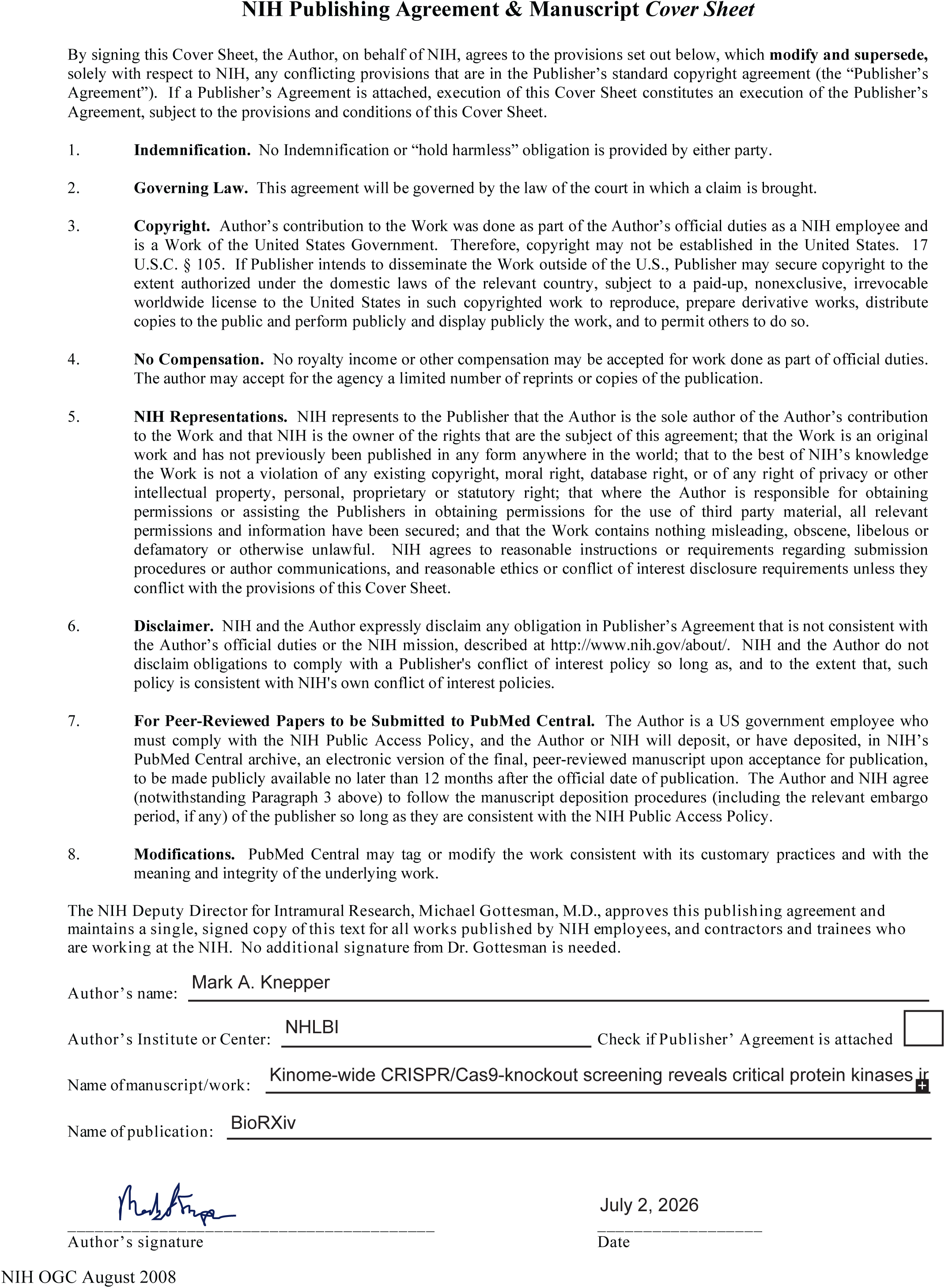

