## Supplemental Figure 1 for "Kinome-wide CRISPR/Cas9-knockout screening reveals critical protein kinases in vasopressin V2-receptor signaling"

**Supplemental Figure 1.** Analysis of data from RNA-sequencing of dDAVP-treated Dyrk1a KO cells (four independent clones versus WT cells (also n=4). Cells were exposed to 0.1 nM dDAVP for 48 hrs. **A.** Top 15 Gene Ontology (GO) terms and KEGG pathways from enrichment analyses of DEGs. Gene Ratio indicates the fraction of input DEGs annotated to the term (Count / total DEGs analyzed). **B.** Heatmap of cell-cycle transcripts from the KEGG Cell cycle pathway showing sample-to-sample variability.

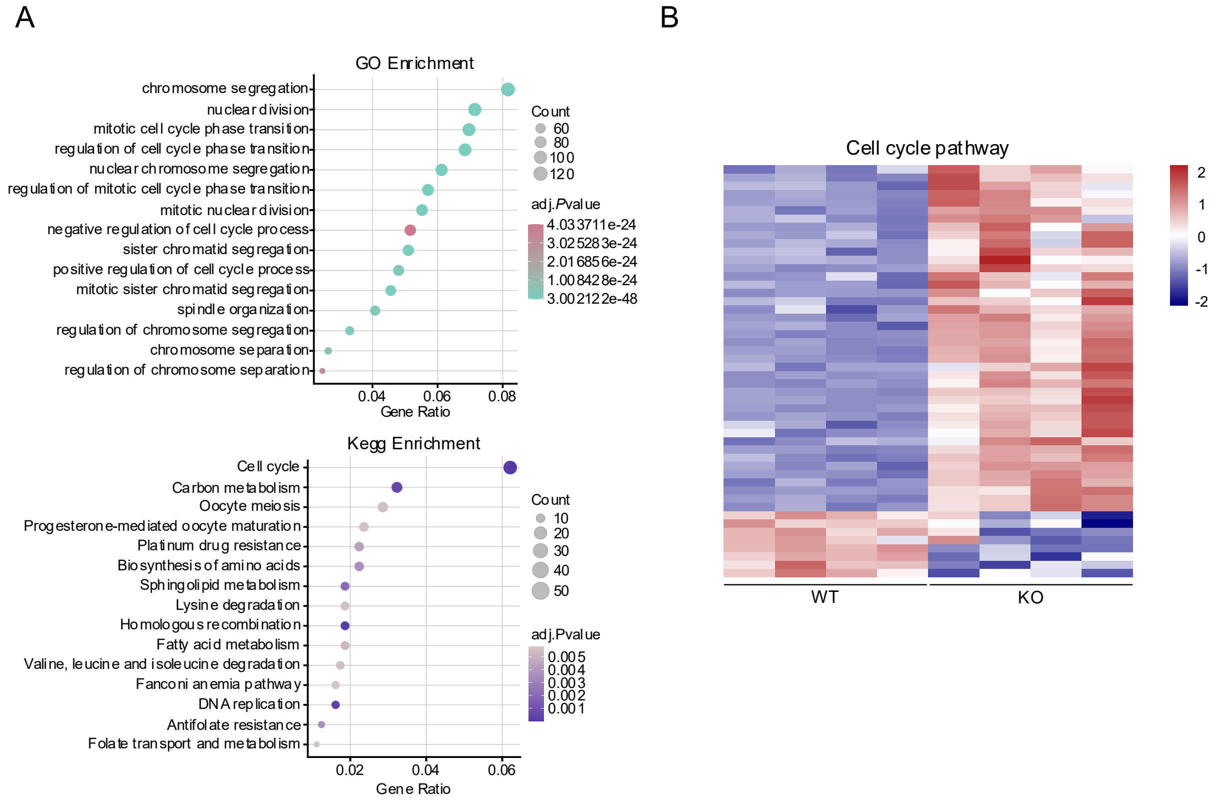
