## Supplementary Table 1 for "Kinome-wide CRISPR/Cas9-knockout screening reveals critical protein kinases in vasopressin V2-receptor signaling"

**Supplementary Table 1. Effect of Stk11 (LKB1) knockout on response to vasopressin analog dDAVP for previously recognized vasopressin-responsive transcripts (RNA-Seq) [*].**

| Gene Symbol* | $\log_2$ (dDAVP/ vehicle), WT | P (t-statistic paired), WT | $\log_2$ (dDAVP/ vehicle), Stk11-KO | P (t-statistic paired), Stk11-KO | Difference, KO - WT |
|---|---|---|---|---|---|
| **Cited1** | 3.60 | **1.38E-04** | 1.01 | **2.00E-02** | -2.59 |
| **Aqp2** | 3.24 | **3.00E-07** | 0.46 | **1.88E-02** | -2.78 |
| **B3gnt7** | 2.28 | **6.36E-07** | 0.85 | **1.40E-02** | -1.42 |
| **Spink8** | 2.04 | **3.36E-06** | 0.25 | 2.09E-01 | -1.79 |
| **C3** | 1.28 | **1.80E-06** | 0.53 | 5.45E-02 | -0.75 |
| **Arl4d** | 1.20 | **3.74E-05** | 0.25 | 5.10E-02 | -0.95 |
| **Selenbp1** | 1.16 | **1.36E-05** | 0.51 | **1.76E-02** | -0.65 |
| **Bcat1** | 1.16 | **5.43E-06** | 0.30 | **1.20E-02** | -0.86 |
| **Sult1c2** | 1.11 | **9.05E-04** | 0.22 | 2.52E-01 | -0.89 |
| **Muc4** | 1.05 | **2.47E-03** | 0.48 | **1.17E-02** | -0.57 |
| **Arhgef3** | 0.98 | **3.29E-04** | 0.34 | **7.55E-03** | -0.64 |
| **Cdk18** | 0.94 | **1.82E-05** | 0.38 | **2.36E-03** | -0.56 |
| **Lbp** | 0.92 | **4.32E-04** | 0.10 | 2.51E-01 | -0.83 |
| **Cyfip2** | 0.86 | **2.92E-03** | 0.35 | 6.22E-02 | -0.51 |
| **Nfkbiz** | 0.82 | **1.77E-04** | 0.20 | **3.14E-03** | -0.62 |
| **Eya2** | 0.82 | **2.95E-03** | 0.28 | **2.56E-02** | -0.53 |
| **Akap12** | 0.81 | **5.71E-05** | 0.30 | **1.49E-02** | -0.51 |
| **Ndrg1** | 0.67 | **1.52E-04** | 0.49 | **1.23E-02** | -0.18 |
| **Prom1** | 0.63 | **6.74E-05** | 0.06 | 4.72E-01 | -0.57 |
| **Idh1** | 0.49 | **5.16E-06** | 0.09 | **4.69E-02** | -0.40 |
| **Tuba4a** | 0.45 | **2.67E-03** | 0.35 | **3.92E-02** | -0.10 |
| **Tbc1d1** | 0.41 | **3.06E-05** | 0.31 | **4.16E-03** | -0.10 |
| **Fads1** | 0.31 | **3.79E-03** | 0.16 | **2.43E-02** | -0.15 |
| **Stat3** | 0.23 | **5.70E-03** | 0.15 | 5.71E-02 | -0.08 |

*** List of known vasopressin-responsive mRNAs from Sandoval et al. (16).**
**KO, knockout; WT, without transformation. Measurements were obtained in 5 distinct KO and WT mpkCCD clones each. P-values less than 0.05 are indicated in bold.**
